# Population history across timescales in an urban archipelago

**DOI:** 10.1101/2025.01.24.633650

**Authors:** Emma K. Howell, Lauren E. Nolfo-Clements, Bret A. Payseur

## Abstract

Contemporary patterns of genetic variation reflect the cumulative history of a population. Population splitting, migration, and changes in population size leave genomic signals that enable their characterization. Existing methods aimed at reconstructing these features of demographic history are often restricted in their temporal resolution, leaving gaps about how basic evolutionary parameters change over time. To illustrate the prospects for extracting insights about dynamic population histories, we turn to a system that has undergone dramatic changes on both geological and contemporary timescales – an urbanized, near-shore archipelago. Using whole genome sequences, we employed both common and novel summaries of variation to infer the demographic history of three populations of endemic white-footed mice (*Peromyscus leucopus*) in Massachusetts’ Boston Harbor. We find informative contrasts among the inferences drawn from these distinct patterns of diversity. While demographic models that fit the joint site frequency spectrum (jSFS) coincide with the known geological history of the Boston Harbor, patterns of linkage disequilibrium reveal collapses in population size on contemporary timescales that are not recovered by our candidate models. Historical migration between populations is also absent from best-fitting models for the jSFS, but rare variants show unusual clustering along the genome within individual mice, a pattern that is reproduced by simulations of recent migration. Together, our findings indicate that these urban archipelago populations have been shaped by both ancient geological processes and recent human influence. More broadly, our study demonstrates that the temporal resolution of demographic history can be extended by examining multiple facets of genomic variation.

**Significance Statement:** Detailed information about a population’s history can be obtained by studying patterns of genomic variation, but these insights can be limited in their temporal scope. Our investigation of white-footed mice in the urban Boston Harbor archipelago demonstrates how combining multiple summaries of genomic variation enables a more complete reconstruction of population history over both the recent and distant past.

## Introduction

The evolutionary processes and events that shape the history of a population leave signatures in patterns of DNA sequence variation. With advances in genome sequencing, statistical methods, and computational approaches, it is now feasible to reconstruct population sizes, migration rates, and split times in any set of populations for which samples can be obtained (Beichman et al. 2018). In addition to quantifying these important evolutionary parameters, demographic inference is a key step toward the goal of using genomic data to characterize natural selection (Pavlidis et al. 2010; Crisci et al. 2013; Lotterhos and Whitlock 2014; Johri et al. 2022).

Existing approaches for demographic inference can be distinguished by the genomic summary used to draw inferences, the statistical framework they employ, or the assumptions they make. An important but often neglected distinction concerns the timescale over which available strategies are best suited to estimate demographic parameters of interest (Nadachowska-Brzyska et al. 2022). This temporal scope is determined by how rapidly a given genomic signal responds to relevant demographic processes and how quickly the signal decays. For example, tracts of identity-by-descent (IBD) are immediately established by recent common ancestry and readily broken up by recombination during subsequent meioses (Browning and Browning 2012; Thompson 2013). As a consequence, long-distance genotype correlations that reflect this coancestry form sensitive indicators of recent history (Palamara et al. 2012; Harris and Nielsen 2013). In contrast, mutation serves as the record keeper of demographic change for analyses based on nucleotide diversity or allele frequencies at unlinked sites. Here, inferences about the recent past are limited by the low rate at which new mutations occur and by the capacity to observe them in a sample (Adams and Hudson 2004; Keinan and Clark 2012; Robinson et al. 2014). Such properties of the sample can also constrain the temporal resolution of approaches based on the sequentially Markovian coalescent (SMC), where the times to most recent common ancestry (TMRCAs) observed for small numbers of genomes reflect population size history over more ancient timescales (Li and Durbin 2011; Schiffels and Durbin 2014). While efforts have been made to broaden temporal resolution by combining different genomic patterns (Boitard et al. 2016; Terhorst et al. 2017), rarely are the temporal limits of any given method tested (but see Boitard et al. 2016). As demographic inference is extended to a wider range of species encompassing a variety of histories, it is important to consider the timescales to which these genomic analyses provide access.

Continental islands are useful settings for investigating how the interplay between historical and contemporary processes shape genomic patterns of variation. Formed by the marine transgressions that followed Pleistocene glacial cycles, the young geological ages of continental islands and their geographic proximity to the mainland offer multiple routes for the establishment of island populations (Losos and Ricklefs 2009). Colonization may be achieved by short-distance dispersal from nearby mainland areas or by vicariance events if species habitation predates island formation (Losos and Ricklefs 2009). For less vagile species, geological history may constrain the demographic history of island populations. Contemporary anthropogenic influence adds a layer of complexity to the population dynamics underlying these island systems, as human transport can overcome otherwise impermeable barriers to dispersal.

The Boston Harbor archipelago presents a prime opportunity to elucidate the contributions of historical and contemporary demography to genomic diversity in island populations. Ranging in size from 0.4 ha to 105 ha, the islands were formed by the glacial retreat and subsequent sea level rise that followed the Last Glacial Maximum (Olmstead Center for Landscape Preservation 2017). Owing to their abundance of natural resources and proximity to the mainland, the islands in Boston Harbor have served a prominent role in the ecological and cultural history of the region (Luedtke and Rosen 1993; Luedtke 1996; Richburg and Patterson 2005). Prior to their incorporation into the Boston Harbor Islands National Recreation Area in 1996, the islands supported diverse public and private endeavors including farms, schools, military forts, and waste management facilities (Olmstead Center for Landscape Preservation 2017). Although there have been efforts to revitalize the natural landscapes of the Boston Harbor (Richburg and Patterson 2005; Olmstead Center for Landscape Preservation 2017), the lasting impact of human development is evident in the high burden of exotic plant species surveyed among the islands (Elliman 2005).

Despite its dynamic history, multiple islands within this “urban archipelago” support stable populations of the white-footed mouse, *Peromyscus leucopus* (Nolfo-Clements and Clements 2015; Nolfo-Clements 2018), a North American endemic that has colonized a wide range of environments (Osgood 1909; Bedford and Hoekstra 2015). *P. leucopus* is an emerging model system for native species that persist in the face of human disturbance (Munshi-South 2012; Yu et al. 2017; Harris and Munshi-South 2017). For example, in the highly urbanized landscape of New York City, *P. leucopus* are ubiquitous in forested fragments and demonstrate an impressive ability to disperse along even the most marginal “green” corridors formed by roadside vegetation, residential areas, and cemeteries (Munshi-South 2012).

In this paper, we take advantage of the unique geological and ecological legacy of the Boston Harbor to elucidate the impact of historical and contemporary demographic processes on genomic variation in populations of *P. leucopus*. Using whole genome sequences from island and mainland populations of mice, we draw inferences from allele frequencies, patterns of linkage disequilibrium, and properties of rare variants. We illustrate how the insights gained from these distinct signals enrich our understanding of demographic history on multiple timescales.

## Results

### Sampling island and mainland sites within the Boston Harbor

The Boston Harbor archipelago is comprised of 34 small islands and peninsulas just east of the city center. Our study focused on two of the inner harbor islands, Peddocks (74.6 ha) and Bumpkin (12.2 ha), and one mainland site on the World’s End peninsula (Figure 1). Using tissue samples collected from wild-caught *P. leucopus*, we constructed short-read whole genome sequence datasets for mice from each of these three locations (n=37 for Bumpkin Island, n=21 for Peddocks Island, and n=19 for mainland World’s End).

**Figure 1.**
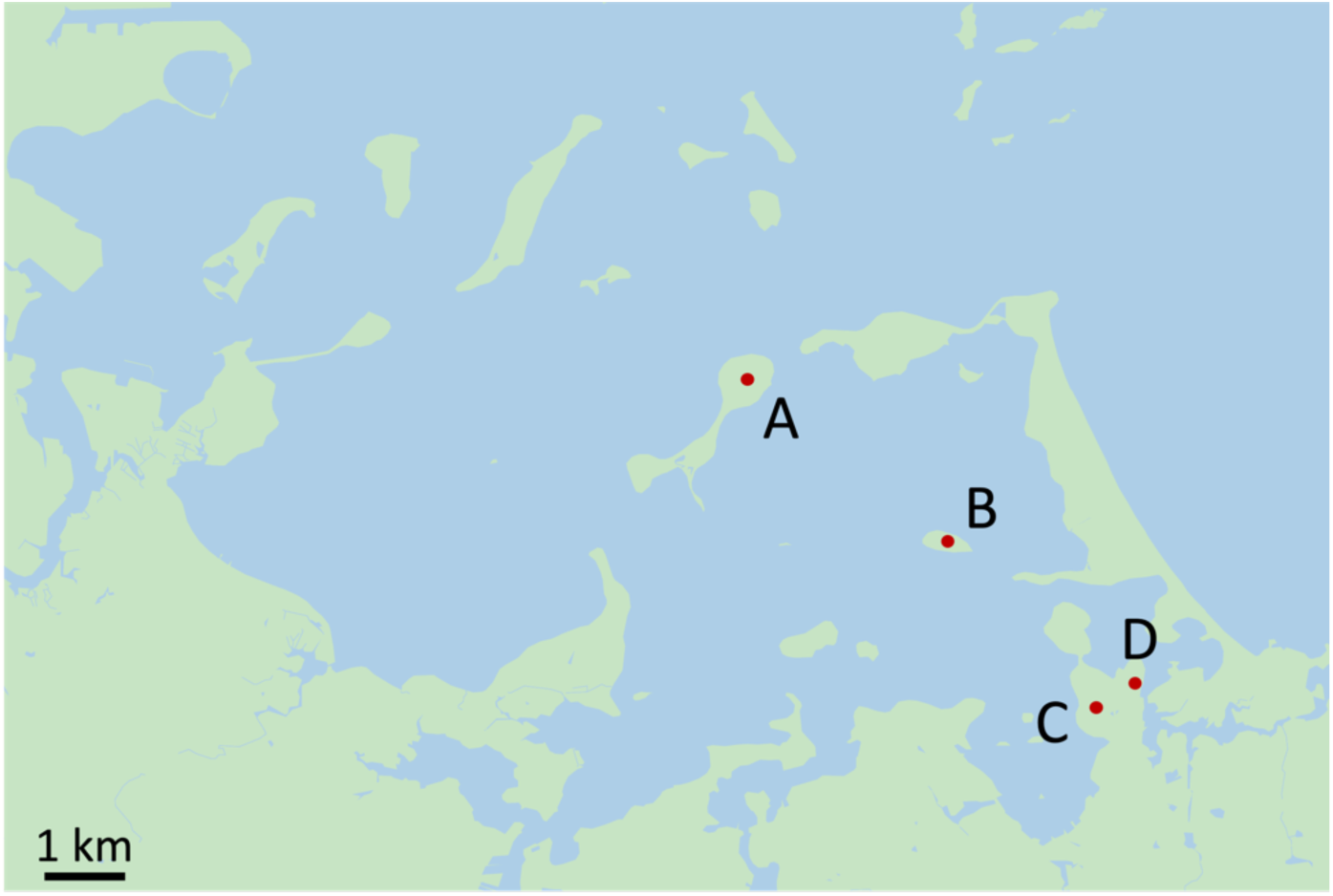
Map of the Boston Harbor archipelago in Boston, Massachusetts. Land areas are depicted in green and water areas are depicted in blue. Red points indicated sampled areas on Peddocks Island (A), Bumpkin Island (B), and mainland World’s End (C and D).

We used a subset of unlinked single nucleotide polymorphisms (SNPs) to estimate pairwise kinship coefficients between sampled mice with KING (Manichaikul et al. 2010). Surprisingly, we inferred a high incidence of close relatives in both island cohorts, with 13.77% of Bumpkin Island pairs and 10.28% of Peddocks Island pairs exceeding the lower bound for 2^nd^ degree relatives (Supplementary figures S1A and S1B). In contrast, only 3.33% of mainland World’s End pairs were inferred as closely related (defined here as pairs ≤ 2^nd^ degree relatives) (Supplementary figure S1C). We did not find any close relatives among mice sampled from different locations.

In addition to this abundance of close relatives, we also identified a single Bumpkin Island mouse (B19) that exhibits substantial genetic divergence from the rest of the Bumpkin sample. Specifically, we found that B19 contributes a disproportionately large number of singleton variants (i.e., those with a minor allele count of 1) to the genome-wide folded site frequency spectrum (SFS) (Supplementary figure S2). Many of these singleton variants are shared with the Peddocks and World’s End cohorts, reducing the likelihood that they originated from sequencing or genotyping errors (Supplementary figure S3). Within-population principal component analyses (PCA) identified this individual as an outlier with respect to the rest of the Bumpkin sample (Supplementary figure S4) and ADMIXTURE (Alexander et al. 2009) analyses conducted on the combined dataset including all three geographic locations suggested that this individual derives ancestry from both World’s End and Peddocks (Supplementary figure S5). Given their potential to bias the SFS, we excluded both B19 and the close relatives we identified in each cohort from allele frequency-based analyses, reducing sample sizes to n=13 for Bumpkin Island, n=13 for Peddocks Island, and n=17 for mainland World’s End.

### Population structure within the Boston Harbor

To characterize genetic relationships between sampled locations, we conducted PCA using genome-wide, unlinked SNPs from each island and mainland cohort (see Materials and Methods). We found that individuals cluster according to sampling location along the first and second principal component axes, indicating genetic divergence between mice inhabiting different sites (Figure 2A). To further assess the degree of structure both within and among sampled locations, we performed model-based inference of global ancestry proportions with ADMIXTURE (Alexander et al. 2009) using the same collection of genome-wide SNPs. Evidence for structure within the combined sample was strong, with cross-validation analyses identifying three ancestral components (Figure 2B) that delineated individuals according to location (Figure 2C). In contrast, a single ancestral component provided the best fit to each of the location-specific samples, suggesting little evidence for structure within each site (Figure 2B). Given these results, we treated each of the sampled locations (Peddocks Island, Bumpkin Island, and mainland World’s End) as representing distinct populations of mice. To orient subsequent demographic analyses, we inspected the SFS of shared and private variants between each pair of populations (Supplementary figure S9). These partitions indicated that mainland World’s End retains much of the genetic variation present in the Peddocks Island and Bumpkin Island samples, providing evidence that the three sampled populations are indeed derived from a recent shared ancestral population.

**Figure 2.**
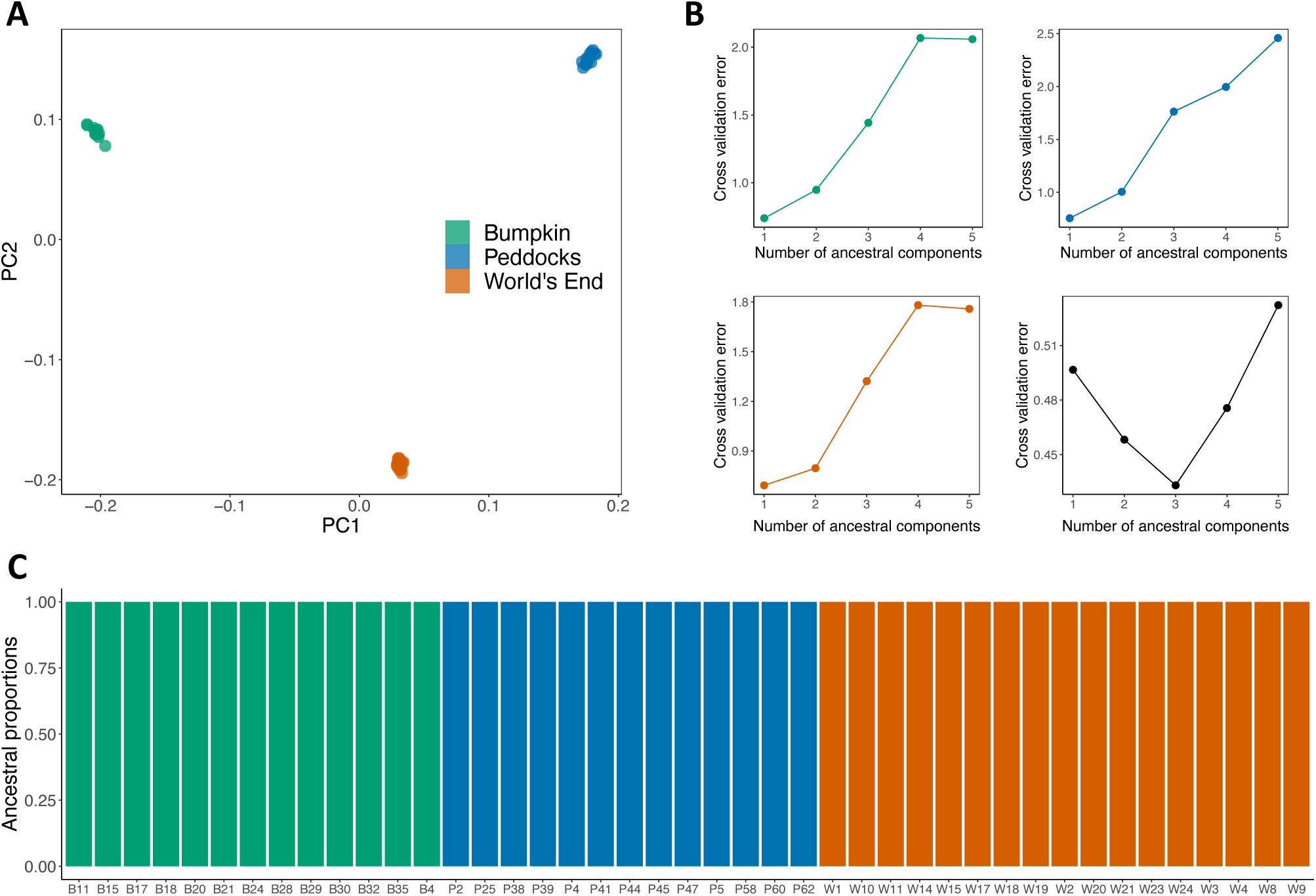
Evidence of population structure among Boston Harbor localities. (A) Principal component analysis conducted using genome-wide, unlinked SNPs. Points represent individuals sampled from either Bumpkin Island (green), Peddocks Island (blue), or mainland World’s End (orange). Percentage of variance explained by PC1=10.36% and by PC2=8.82%. (B) Five-fold cross validation analysis with ADMIXTURE (Alexander et al. 2009). A single ancestral component (k=1) yields the minimum cross validation error within each of the sampled localities (top left, top right, and bottom left). In the combined sample (bottom right), three ancestral components (k=3) yield the minimum cross validation error. (C) Global ancestry proportions among sampled individuals, assuming k=3 ancestral components. Each individual is represented by a colored bar. The height of each colored segment in a bar gives the proportion of ancestry an individual derives from a given component. Ancestral component colors reflect population of origin (as in Figure 2A).

### Historical insights about demography

To reconstruct the demographic histories of these island and mainland populations, we employed an approach based on the joint site frequency spectrum (jSFS). Using *Moments* (Jouganous et al. 2017), we fit two-population demographic models to the folded jSFS of putatively neutral variants observed in each pair of population samples (see Materials and Methods). Our aims were to estimate population divergence times, changes in effective population size (N_e_) through time, and historical levels of gene flow between populations. For each pair of populations, our model selection criteria (see Materials and Methods) favored the simplest model of population divergence. This model involves a split between sampled populations from a shared ancestor, constant island and mainland N_e_ following their split, and no migration (Supplementary figure S6A). Maximum likelihood estimates of N_e_ and divergence times yielded congruent values across the three pairwise comparisons (Figure 3). Parameter estimates across these three inferred histories suggest a large ancestral N_e_ (ranging from N_e_=413,589 to N_e_=470,513), small post-split N_e_ for both Bumpkin Island (between N_e_=7,257 and N_e_=7,501) and Peddocks Island (between N_e_= 9,465 and N_e_= 9,669), and a larger post-split N_e_ for mainland World’s End (between N_e_= 65,483 and N_e_=67,573) (Figure 3). Although there is overlap among the confidence intervals for all inferred divergence times, their point estimates imply an older split between the Peddocks and World’s End populations (7,659 generations ago) and a more recent split between the Bumpkin and World’s End populations (6,911 generations ago) (Figure 3).

**Figure 3.**
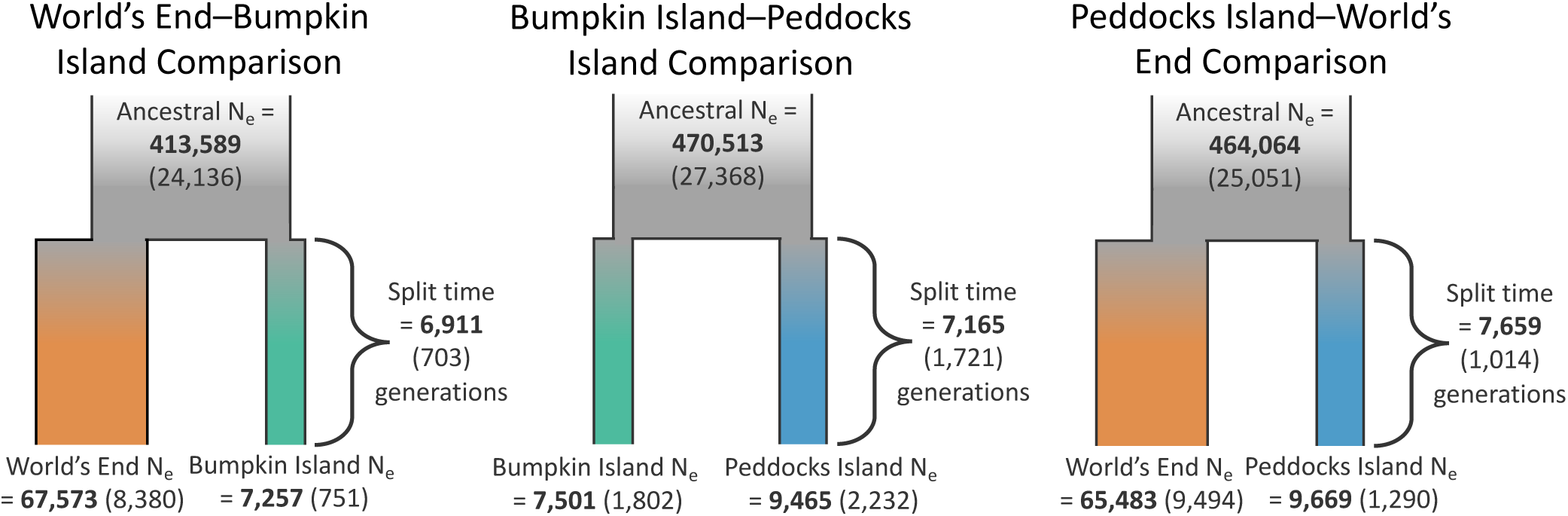
Demographic inferences from the SFS support old divergence times and smaller N_e_ for island populations. Model fitting and parameter estimation were performed separately for the jSFS constructed from each pair of populations, yielding three separate histories for the mainland World’s End-Bumpkin Island (left), Bumpkin Island-Peddocks Island (middle), and Peddocks Island-mainland World’s End (right) comparisons. Maximum likelihood estimates for each parameter are bolded. Standard deviations for each parameter estimate are given in parentheses and were ascertained using the Godambe Information Matrix (Coffman et al. 2016) as an alternative to conventional bootstrapping.

To evaluate the ability of the inferred demographic histories to recover summaries of genetic variation beyond the jSFS, we conducted predictive simulations under the best-fit parameter estimates using *msprime* (Baumdicker et al. 2022) (see Materials and Methods). For π, Tajima’s *D*, and F_ST_, we found close agreement between the means of the observed window-based estimates and simulated distributions (Figure 4). For all three measures, the models underpredicted the variance in the observed summary statistic distributions (Figure 4), but much of this discrepancy is likely driven by unmodeled heterogeneity in the recombination and the mutation rate as well as differences in SNP density caused by the filtering conducted on the empirical callset. In contrast to the close fit our inferred model provides to allele frequency-related summaries, we found that linkage disequilibrium (LD) decays more quickly and reaches a lower minimum at 1 Mbp for simulated data (Figure 4). This discrepancy is largest for the two island populations, Bumpkin and Peddocks, where LD decay curves are unusually high and flat. In both cases, r^2^ exceeds 0.25 even for SNPs separated by as much as 1 Mbp (Figure 4).

**Figure 4.**
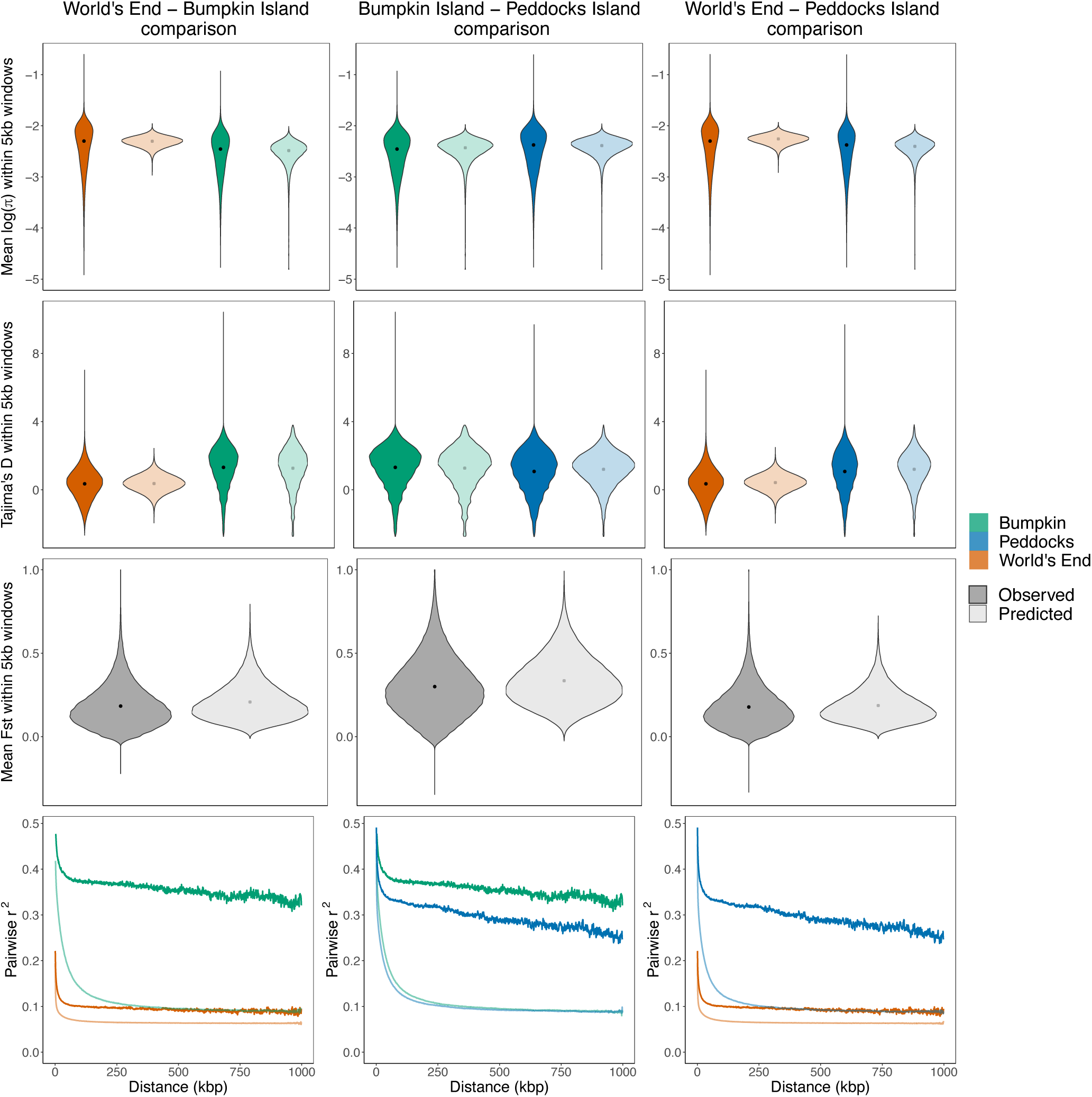
SFS-based demographic models provide a close fit to some, but not all, summaries of variation. Panels compare summaries of variation measured from empirical data (dark shading) to those obtained from predictive simulations conducted using best-fit demographic parameter estimates (light shading). Violins plot the distributions of π (log-scaled; first row), Tajima’s D (second row), and F_ST_ (third row) computed in 5 kbp windows. Black points denote the mean value of each distribution. LD decay curves (fourth row) plot average pairwise r^2^ between SNPs separated by increasing physical distances (up to 1 Mbp). For single-population summaries, empirical and simulated distributions/curves are colored according to population (Bumpkin Island, green; Peddocks Island, blue; mainland World’s End, orange). Columns titles denote the demographic model used for each set of predictive simulations (World’s End-Bumpkin Island, left; Bumpkin Island-Peddocks Island, middle; World’s End-Peddocks Island, right).

Beyond these common summaries of genomic variation, we detected an additional genomic pattern that our simulations failed to predict. In both the Bumpkin Island and Peddocks Island samples, singleton variants form distinct tracts in which a single mouse contributes a disproportionate number of singletons to the local (within-population) SFS (Figures 5A and 5C). At the genomic scale, this pattern results in substantial inter-chromosomal variance in individual singleton contributions (Supplementary figures S10 and S11), with multiple Bumpkin mice deriving more than 50% of their contributed singletons from a single chromosome (Supplementary figure S10). This pattern is less evident in mainland World’s End, where singleton variants appear more uniformly distributed across the genome (Figure 5E) and across mice (Supplementary figure S12). Chromosome-scale predictive simulations demonstrate that the extreme spatial clustering of singletons observed in the island samples is unlikely to be produced solely by the inferred reductions in population size (Figures 5B, 5D, and 5F).

**Figure 5.**
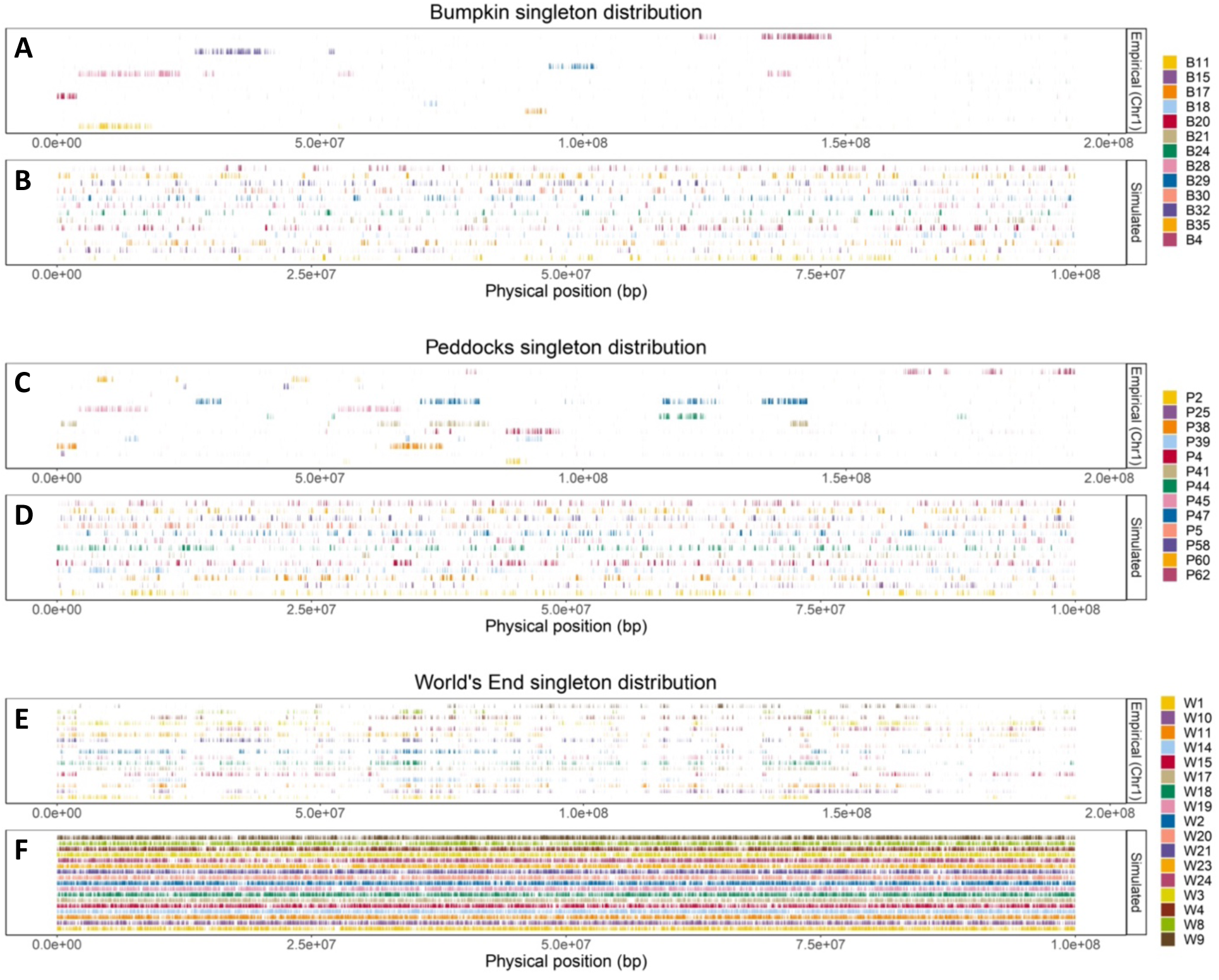
SFS-based demographic models fail to recover the spatial clustering of rare variants observed in island samples. (A, C and E) Distribution of singleton variants observed on chromosome 1 (NC_051063.1) within the Bumpkin Island (A), Peddocks Island (C), and mainland World’s End (E) samples. X-axes give the physical position along chromosome 1. Vertical, colored tick marks denote observed singleton variants. Each row reflects the singleton variants of a distinct individual. (B, D, and F) Corresponding singleton distributions predicted by the inferred demographic histories for Bumpkin Island (B), Peddocks Island (D), and mainland World’s End (F). X-axes give the location of singletons along a 100 Mbp simulated chromosome and tick marks denote singleton variants observed in each simulated individual. For clarity, we only show the simulated singleton maps derived from the Bumpkin-World’s End demographic model (for B and F) and the Peddocks-World’s End demographic model (for D), though we note that patterns are similar regardless of which specific demographic model is used for a given population.

In the following sections, we explore possible causes for these intriguing features of the data that are not fit by our historical models of demography based on the jSFS.

### Inter-archipelago migration may explain the observed clustering of rare variants

The unusual distribution of singleton variants in the island samples may be a more localized manifestation of the excess singletons we observed in the excluded Bumpkin Island mouse, B19 (Supplementary figure S3). Our finding that B19’s singletons are segregating in both the Peddocks and World’s End populations (Supplementary figure S3) raises the possibility that this pattern is driven by gene flow within the archipelago. Indeed, the proximity of island and mainland areas within the Boston Harbor, combined with the frequent human traffic between them, leaves multiple routes for the migration of small mammals. Given the substantial genetic divergence we observed between the three populations (Figure 2), we should expect much of the genomic content exchanged by within-archipelago migration to involve private variation.

Consequently, when tracts of migrant ancestry are observed in the recipient population, the private variation they carry will manifest as new polymorphisms. If migration is sufficiently rare or recent such that individuals carry distinct migrant haplotypes, then these tracts of migrant ancestry will contribute disproportionally to rare allele frequency bins.

We used coalescent simulations of genome-scale datasets to test this verbal model and explore the possibility that migration (absent from our candidate demographic histories) could explain the pattern of singleton clustering observed in both island samples. For each pair of populations, we augmented our best-fit demographic model with migration from an unsampled “ghost” population, varying the migration rate and length of time during which migration occurs (see Materials and Methods).

By inspecting the spatial distribution of singleton variants across simulated chromosomes, we found qualitative evidence that migration can generate dense tracts of singletons, such as those observed in both the Bumpkin Island and Peddocks Island samples (Supplementary figures S14 and S15, respectively). Specifically, regimes involving either extended periods of weak migration (Figures 6A and 6C) or brief periods of strong migration (Figures 6B and 6D) yield singleton tracts that are similar in length and abundance to those found in the island datasets (Figures 5A and 5C).

**Figure 6.**
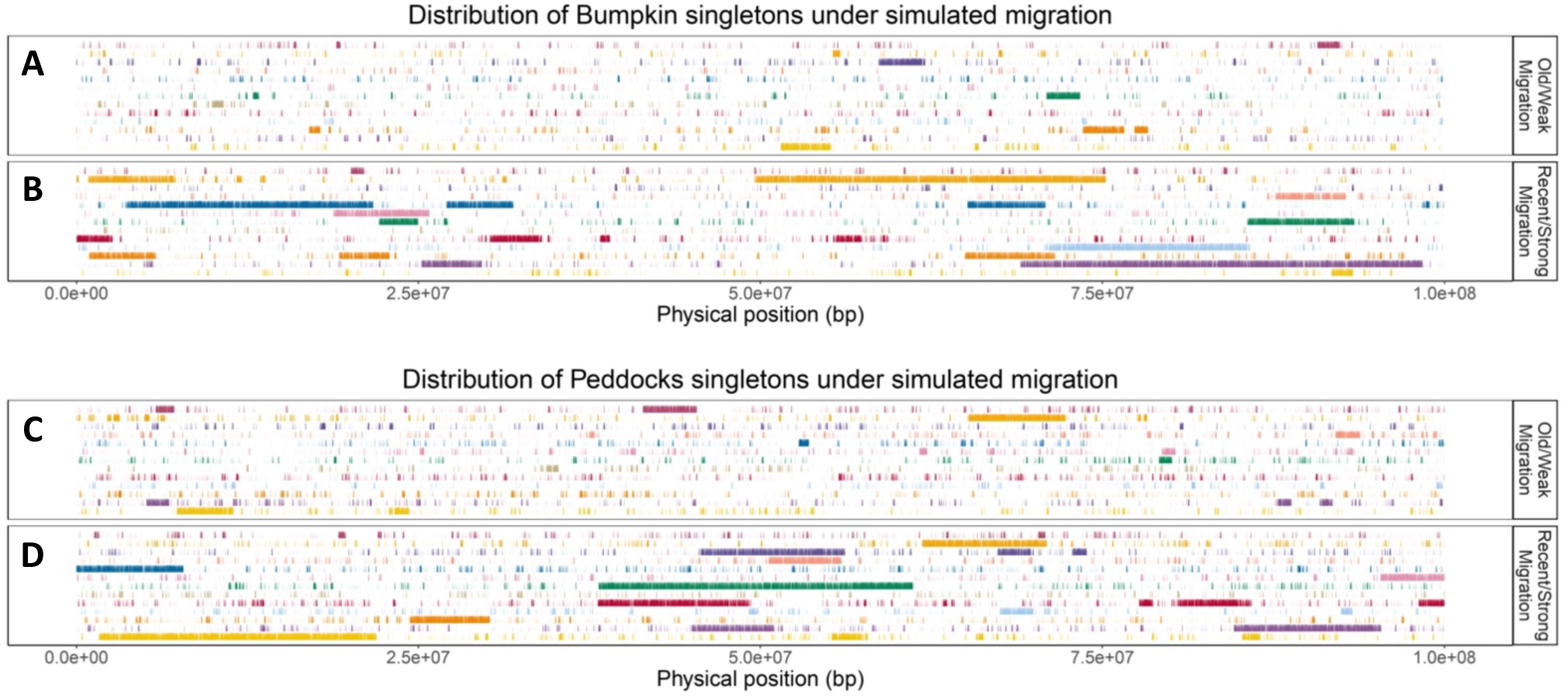
The physical clustering of rare variants is recovered by simulated migration regimes. (A and C) Singleton distributions predicted for Bumpkin Island (A) and Peddocks Island (C) samples under a demographic history that includes weak migration (at a rate of 1e-5) between sampled populations and an unsampled “ghost” population that occurs for 10% of the total divergence time. (B and D) Predicted singleton distributions for Bumpkin (B) and Peddocks (D) samples under a demographic history that features stronger migration (at a rate of 1e-3) occurring for only 1% of the total divergence time. Singleton maps are arranged as in Figure 5B and 5D. For clarity, we only show the simulated singleton maps derived using the population size and divergence time estimates from the Bumpkin-World’s End (A and B) and Peddocks-World’s End (C and D) demographic models, though we note that patterns are similar regardless of which specific demographic model is used as the basis for each set of migration simulations.

The other salient feature of singleton variants in the island samples is their uneven distribution among mice. We found high variance in the number of singletons each mouse contributes to the SFS of a given chromosome (Supplementary figures S10 and S11). To evaluate whether the simulated migration regimes could recover this variability, we compared chi-squared test statistics computed for each empirical and simulated chromosome assuming a null hypothesis of equal singleton contributions among mice. We found that the mean chi-squared statistic measured across empirical Peddocks Island and Bumpkin Island chromosomes falls within the range of values produced by the simulated migration regimes (Supplementary figure S13A-B). Although these empirical chi-squared test statistics are more extreme than those obtained under simulations without migration, even migration-free simulations yield large deviations from the null hypothesis. These results indicate that the population size history represented in our candidate demographic models may itself affect the distribution of singleton variants among individuals.

To develop intuition for how population size history and migration interact to shape individual singleton contributions, we leveraged the functionality of the tskit library (Kelleher et al. 2018) to inspect properties of the local genealogies underlying simulated chromosomes (see Materials and Methods). Focusing on Bumpkin Island chromosomes simulated under the Recent/Strong migration regime featured in Figure 6B, we compared various measures of branch lengths between genealogical trees spanning regions with and without migrant ancestry (Figures 7A and 7B). As expected, the presence of one or more migrant haplotypes increases the total external branch length (*E*), the mean external branch length within a tree (Mean(*E*)), and the standard deviation of external branch lengths within a tree (Sd(*E*)) (Figure 7C). Interestingly, despite the large distortions it produces along external branches, migrant ancestry appears to have little effect on the total tree height (*H*) or the total branch length (*T*) (Figure 7C). In addition, 100% of trees spanning migrant haplotypes and 99.99% of trees without migrant haplotypes find their most recent common ancestor (MRCA) in the ancestral population (as opposed to the focal island or ghost population). This congruency in the timing and source of the MRCA between tree categories suggests that the private variation carried on migrant haplotypes reflects a subset of ancestral variation that has been lost in the focal population rather than new mutations that occurred within the source population.

**Figure 7.**
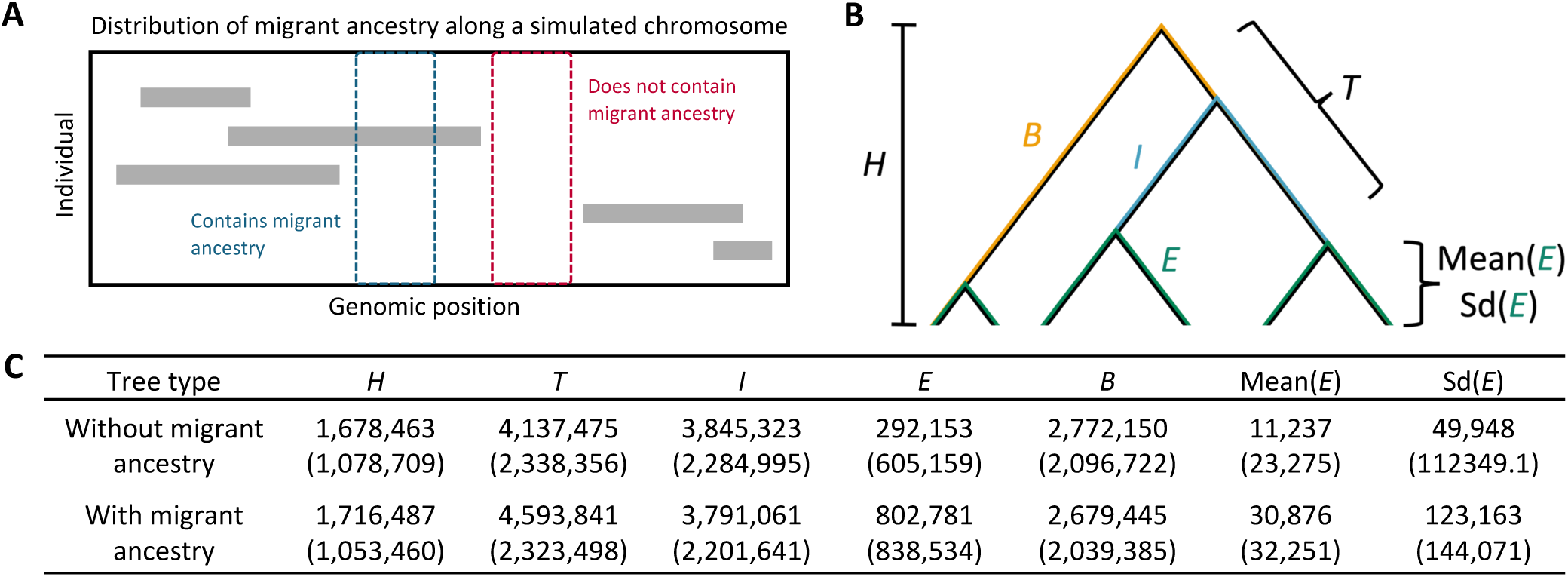
Singleton clustering reflects the interaction of migration and population size history on the external branch lengths of local genealogies. We inspected coalescent genealogies for the Bumpkin Island sample simulated under the Recent/Strong migration regime featured in Figure 6B. (A) Diagram illustrates how local genealogies were classified into two types. Each row represents variation observed in a different simulated individual. Gray rectangles reflect migrant haplotypes derived from the “ghost” population. The tskit library (Kelleher et al. 2018) was used to distinguish between local genealogies spanning regions containing one or more migrant haplotypes (blue box) and those spanning regions without migrant ancestry (red box). (B) We measured multiple genealogical quantities for each category of trees: the total tree height (*H*), sum of total branch lengths (*T*), sum of basal branch lengths (*B*), sum of internal branch lengths (*I*), sum of external branch lengths (*E*), mean external branch length within a tree (Mean(*E*)), and standard deviation of external branch lengths within a tree (Sd(*E*)). (C) Table summarizes the mean value of each quantity across all trees observed along a simulated 100 Mbp chromosome. Standard deviations across trees are given in parentheses.

### Small contemporary effective population sizes may explain elevated linkage disequilibrium

The pattern of LD decay we observed in the Bumpkin Island and Peddocks Island populations yielded curves that are unusually high and flat. Such characteristics may indicate a recent and severe population collapse (Rogers 2014). In addition to this trend of elevated LD, the abundance of close relatives we identified in each sample may signify smaller effective population sizes than suggested by our candidate demographic models (Wang 2009; Waples and Anderson 2017).

Given these findings and the dynamic ecological histories of the islands, we hypothesized that the demographic models we inferred from the jSFS may not capture more recent changes in effective population size. To investigate this possibility, we reconstructed contemporary N_e_ using two approaches that leverage different facets of LD (see Materials and Methods). *CurrentNe* uses LD measured between unlinked markers to obtain a point estimate of N_e_ that reflects the several most recent generations (Santiago et al. 2024). In contrast, GONE uses LD measured between markers separated by increasing genetic distances to reconstruct the trajectory of N_e_ in the recent past (Santiago et al. 2020). Using these approaches, we estimated recent N_e_ and the recent N_e_ trajectory for each population using either the full sample or the subset of unrelated individuals (excluding relationships ≤ 2^nd^ degree).

Regardless of whether the sample included close relatives, point estimates of contemporary N_e_ for each population are over an order of magnitude smaller than suggested by best-fit demographic models (Figure 8A and Figure 3). For the unrelated samples of mice, we obtained N_e_ point estimates of 20, 66, and 87 for the Bumpkin Island, Peddocks Island, and mainland World’s End populations, respectively (Figure 8A). As expected, datasets that were pruned for close relatives yielded larger N_e_ estimates (Figure 8A). Recent N_e_ trajectories inferred with GONE are similar between datasets with and without close relatives, with the exception of the recent bottleneck inferred for Peddocks from the full sample (Figure 8B). For all populations, N_e_ estimates in the immediate past (generation 0) are close to the point estimates obtained with *currentNe* (Figures 8A and 8B). These trajectories suggest that each population has experienced dynamic changes in N_e_ over the last 150 generations (Figure 8B).

**Figure 8.**
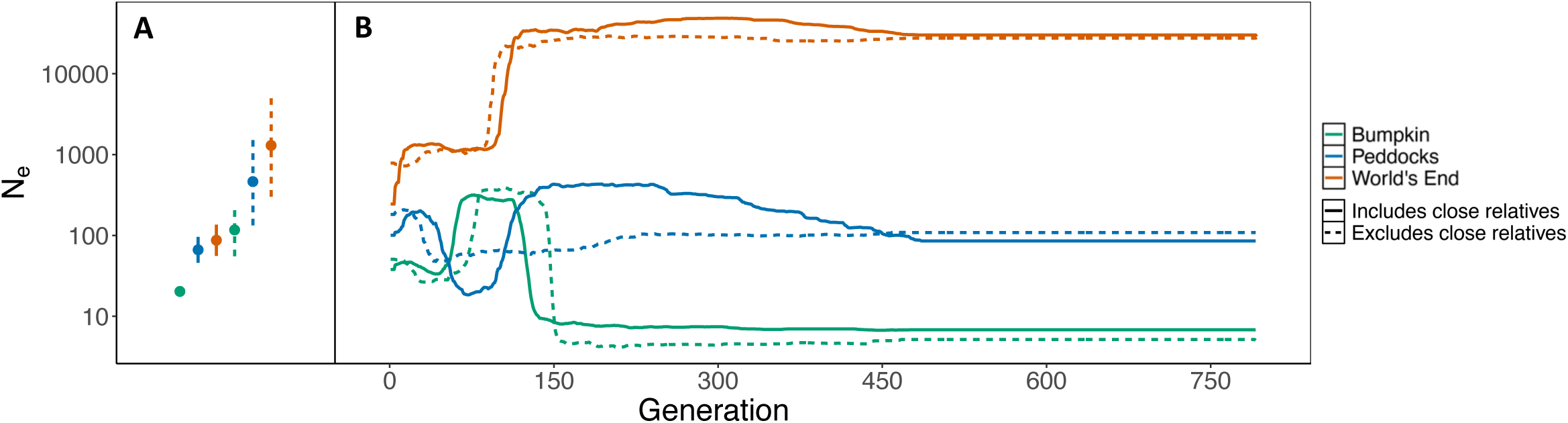
LD-based estimates of recent effective population size are an order of magnitude smaller than SFS-based inferences. (A) Point estimates of contemporary effective population size obtained with *currentNe* (Santiago et al. 2024). Vertical lines represent 90% confidence intervals. (B) Recent effective population size trajectories estimated by GONE (Santiago et al. 2020). N_e_ trajectories span the present (generation 0) to 750 generations into the past. Point estimates and N_e_ trajectories are colored according to population (Bumpkin Island, green; Peddocks Island, blue; mainland World’s End, orange). Dotted confidence intervals/trajectories reflect estimates made using a subset of the data in which close relatives (defined as relationships closer than or equal to 2^nd^ degree) were excluded. Solid confidence intervals/trajectories reflect estimates made using the full sample.

## Discussion

Demographic histories of natural populations are shaped by both historical processes (e.g., glacial cycles, sea-level changes, and tectonic activity) and contemporary shifts (e.g., anthropogenic changes in climate, habitat loss, and fragmentation). While fully capturing the dynamic nature of these histories is beyond the capacity of population genetics, increasingly complex insights about population history are being extracted from genomic data. Despite this progress, we lack a unified framework for drawing inferences about demographic history that span the recent and distant past (Nadachowska-Brzyska et al. 2022). Using island and mainland populations of endemic mice as a model system, we illustrate the importance of timescale in the interpretation of empirical patterns of genomic variation. In the following sections, we discuss both the historical and contemporary insights gained about *P. leucopus* in the Boston Harbor and conclude with general prospects for conducting demographic inference along the temporal continuum.

### Historical insights

The candidate demographic models we obtained from our SFS-based inference suggest that the current distribution of *P. leucopus* in the Boston Harbor was primarily mediated by the geological formation of the archipelago. The topography of the Boston Harbor was shaped by Pleistocene glacial activity which formed the drumlin field that constitutes islands within the archipelago (Kaye 1982; Aubrey 1996). Rises in sea level and the subsequent inundation of the Boston Harbor basin severed existing land connections and established the present-day distribution of island and mainland areas (Kaye 1982; Aubrey 1996). Historical shoreline reconstructions suggest that the islands were isolated 3,000-6,000 years ago (Aubrey 1996).

Assuming 1-2 generations per year (as observed in natural populations of *P. leucopus* in Michigan (Burt 1940)) our inferred divergence times, ranging from 6,000 to 7,000 generations ago, show broad agreement with the estimated timing of island formation. While overlapping confidence intervals raise uncertainty about the order of inferred population splits, their point estimates are consistent with the piecemeal changes in connectivity suggested by the geological history of the region, in which outer islands such as Peddocks were isolated earlier from the mainland (and are therefore older) than the inner islands of the archipelago such as Bumpkin (Aubrey 1996). Although such comparisons should be interpreted cautiously in light of uncertainty about generation time and mutation rate in these populations, our inferences about divergence times are largely congruent with the geological formation of the Boston Harbor archipelago.

Demographic models based on the SFS also indicate that all three populations experienced reductions in N_e_ following their divergence from a shared ancestor. These size reductions were most extreme for the populations on Bumpkin (60-fold decrease) and Peddocks (49-fold decrease). The modest N_e_ values inferred for the island populations could reflect founder effect bottlenecks that accompanied their isolation from the mainland, or constraints on habitat and resource availability driven by the small land areas of both Bumpkin (12.2 ha) and Peddocks (74.6 ha).

These historical insights provide useful context for understanding phenotypic evolution in this system. Mice surveyed on both Bumpkin and Peddocks are 40-50% heavier than mainland *P. leucopus* (Nolfo-Clements et al. 2017). This extreme phenotypic difference suggests that mice within the Boston Harbor follow the “island rule”, a trend in which large-bodied species become smaller on islands and small-bodied species become larger (Foster 1964; Van Valen 1973; Lomolino 1985; Benitez-Lopez et al. 2021). The vicariance mechanism of island colonization suggested by our inferred histories provides a contrast to classic examples of the island rule in house mice that involve recent, human-mediated introductions (Berry 1964; Berry et al. 1978a; Berry et al. 1978b; Berry et al. 1979; Rowe-Rowe and Crafford 1992). Unlike the rapid evolution of body size observed in these cases, our findings raise the possibility that mice in the Boston Harbor evolved large bodies over a longer period of time.

### Contemporary insights

The divergence times and historical N_e_ values estimated from the SFS provide a useful starting point for reconstructing the demographic history of *P. leucopus* in the Boston Harbor. By employing orthogonal approaches to N_e_ estimation and by leveraging the unique empirical pattern we observed among singleton variants in the island populations, we find strong evidence for a more dynamic recent history than indicated by this candidate model. Our results suggest an important role for recent human activity in shaping multiple demographic parameters.

The stark contrast between the LD decay predicted by our candidate models and the LD decay observed in the empirical data may be explained by the differences in N_e_ that we estimated using SFS-based versus LD-based approaches. Although the rank order of N_e_ among island and mainland populations is consistent across both historical and contemporary estimates, their magnitudes differ considerably. Multiple studies have found that empirical patterns of LD are particularly difficult to recover with models derived from the SFS (Harris and Nielsen 2013; Garud et al. 2015; Beichman et al. 2017), but the source of this discrepancy is rarely investigated. In *P. leucopus* from the Boston Harbor, patterns of LD are consistent with very small N_e_ in the recent past, with variation in the temporal trajectory of N_e_ among populations starting at 150 generations ago. These findings could reflect the ecological transformation of the Boston Harbor during the last 400 years. European colonization initiated dramatic shifts in the natural landscape through the clear-cutting of forested islands for wood and the conversion of land into sites for agricultural, military, and public use (Richburg and Patterson 2005; Olmstead Center for Landscape Preservation 2017). This intense human usage has likely decreased habitat and resource availability for native species like *P. leucopus*.

Regardless of the cause for small contemporary N_e_, our findings have important implications for the future of these populations. Small N_e_ limits the amount of additive genetic variance that can be maintained, increasing the likelihood of extinction in the face of environmental change (Lande and Barrowclough 1987; Bürger and Lynch 1995; Kardos et al. 2021). Even in the absence of new or changing selective pressures, the continual input of deleterious mutations leave small N_e_ populations vulnerable to extinction by “mutational meltdown”, a compounding sequence of events in which the fixation of deleterious mutations leads to declines in population size and subsequent increases in the rate at which additional deleterious mutations are fixed (Lynch et al. 1995).

The recent history of *P. leucopus* populations in the Boston Harbor probably featured changes in migration, in addition to dynamic population sizes. Our simulations of ghost migration indicate that inter-archipelago migration is a likely (though not definitive) cause for the unusual partitioning of singleton variants in the island mice. Despite the geographic proximity of island and mainland sites in the Boston Harbor, the ocean could impose a strong barrier to the natural dispersal of *P. leucopus*. However, portions of the harbor occasionally freeze, allowing for mammals to move between island and mainland areas by crossing the ice (Nolfo-Clements 2018). In addition, Bumpkin Island experiences transient connections to the mainland via a land bridge during periods of low tide (Nolfo-Clements and Clements 2015). A more general mechanism for the migration of *P. leucopus* may be their inadvertent transport by humans. The Boston Harbor is a popular recreation area and both public ferries and private watercraft generate substantial island-island and island-mainland traffic.

To better understand why our candidate SFS-based demographic models failed to support a history with migration, we explored how ghost migration manifests in the jSFS used for inference (see Materials and Methods). Under our simulated scenarios, the signal that ghost migration leaves in the jSFS derives from both the movement of shared ancestral polymorphisms and the introduction of variants that are private to the unsampled population. The strength of this signal depends on the rate of migration and the duration over which it occurs (Supplementary figures S16 and S17). Comparisons of the island-mainland jSFS under models with and without ghost migration reveal that only high intensity regimes (i.e., those involving a high rate of migration over a long period of time) can produce correlations in the island-mainland jSFS that extend to shared intermediate allele frequencies (Supplementary figures S16 and S17). Notably, the types of migration capable of producing the observed singleton clustering (Figure 6) have either a negligible impact on the jSFS (for the Old/Weak regime), or one that is restricted to private and low allele frequencies (for the Recent/Strong regime) (Supplementary figures S16 and S17). These findings raise the possibility that the migration experienced by the island populations occupies a zone of parameter space that precludes robust estimation by SFS-based approaches but is nonetheless sufficient to create the observed pattern of singleton clustering.

### Prospects for demographic inference along the temporal continuum

As demographic inference is increasingly aimed at characterizing the histories of endangered, fragmented, and declining populations (Morin et al. 2020; Beichman et al. 2021; Kyriazis et al. 2024), it is important to jointly consider estimates of N_e_ that capture historical levels of variability and those that reflect the immediate past. Approaches for estimating contemporary N_e_ are widespread in conservation genetics (Luikart et al. 2010) but have seen limited application in the broader field of population genetics. Yet, these estimates may be particularly relevant for understanding processes that occur over ecological timescales, such as seasonal adaptation in *D. melanogaster* (Bergland et al. 2014; Machado et al. 2021). They may also be important for forecasting the future evolutionary trajectories of populations, especially in the face of anthropogenic change.

In addition to motivating deeper consideration of N_e_, our investigation suggests new avenues for reconstructing contemporary migration histories. We find that the distributions of rare variants along the genome and across sampled individuals serve as sensitive indicators of recent or rare gene flow. In contrast to SFS-based approaches, which treat SNPs in different genomic locations as interchangeable (Sawyer and Hartl 1992; Gutenkunst et al. 2009), the signature of recent migration we discovered leverages the positions of SNPs along chromosomes. Furthermore, detecting this signature requires no knowledge or sampling of potential source populations.

Few approaches exist to drawing inferences about migration from an unsampled ghost population. A commonly used statistic, *S*,* relies on the capacity for introgressed ancestry to partition a sample in a congruent manner over long genomic distances (Wall 2000; Plagnol and Wall 2006). The singleton pattern we describe appears suited to detect migration in cases where it is too weak to generate such patterns of LD. Future work is necessary to characterize the dependence of this signal on levels of ancestral diversity, divergence times between populations, and sample size. More detailed inferences may be achieved by extending existing theory about the distribution of external branch lengths to incorporate gene flow and population size change (Blum and François 2005; Caliebe et al. 2007; Disanto and Wiehe 2020).

Altogether, our study illustrates how combining multiple features of genomic variation can improve the temporal breadth of demographic inference. While substantial progress remains to be made in bridging the gap between the recent and distant past, important insights can still be gained by deeper evaluation of model fit and interrogation of empirical patterns.

## Materials and Methods

### Population sampling and sequencing

#### Animal trapping

Animals were captured using Sherman live traps spaced at approximately 7 m intervals in grids or transect lines depending on the location. Traps were baited with a mixture of peanut butter and oats. Bedding in the form of dried leaves was added to the traps to assist with thermoregulation. We monitored the traps on Bumpkin for a total of 380 trap nights, for 480 trap nights on Peddocks, and 200 trap nights on World’s End. Trapping for this study occurred June-August 2019.

#### Animal handling and sampling

We checked traps once daily during trapping intervals. Trapped individuals were transferred into a large, unsealed plastic bag to allow for species identification, sexing, maturity evaluation, and weighing with a spring scale. Adult and young adult mice were sampled using a 2 mm tissue punch applied to the right ear. Animals were subsequently released. We chose this sampling method due to its low level of invasiveness and because many captured wild mice have similar injuries to their ears that we presume have occurred naturally. Tissue samples were preserved and transported in 95% ethanol-filled polypropylene microcentrifuge tubes and chilled on ice. The ear punch and forceps were sterilized in between samplings using 95% ethanol to prevent cross contamination. This work was approved by the National Parks Service’s Institutional Animal Care and Use Committee (project NER_BOHA_Nolfo-Clements_SmMammals_2019.A3).

#### Whole genome sequencing

DNA was extracted from ear punch tissue samples using Gentra Puregene Tissue Kits (Qiagen, Germantown, MD, USA). The purity of extracted DNA was assessed by measuring 260/280 absorbance ratios with a NanoDrop Spectrophotometer (Thermo Fisher Scientific, Waltham, MA, USA) and sample concentrations were measured using the Qubit dsDNA HS Assay (Life Technologies, Carlsbad, CA, USA). Prior to library preparation, the presence of high-molecular weight DNA was confirmed using agarose gel electrophoresis. Library preparation and sequencing were conducted by the University of Wisconsin-Madison Biotechnology Center. Sequencing libraries were constructed with the Illumina DNA Library Prep using 100-500 ng of input DNA per sample. Purified and pooled libraries were sequenced on the Illumina NovaSeq6000 platform to generate paired-end 150 bp reads. Samples were sequenced to a targeted coverage of 30X per individual.

### Sequence data processing and variant calling

#### Sequence alignment

Raw sequencing reads were pre-processed to remove adapter sequences and low-quality base calls. Illumina universal adapter sequences were removed using BBDuk v38.96 (https://sourceforge.net/projects/bbmap/) and low-quality base calls were trimmed with Trimmomatic v0.39 (Bolger et al. 2014). Processed sequencing reads were then aligned to the *P. leucopus* reference genome (UCI_PerLeu_2.1; GenBank accession: GCA_004664715.2; Long et al. 2019) using BWA-MEM v0.7.17 (Li and Durbin 2009).

#### Variant calling and filtering

We adapted our variant calling approach from the Genome Analysis Toolkit (GATK) Best Practices recommendations for germline short variant discovery (McKenna et al. 2010; Van der Auwera and O’Connor 2020). Duplicate reads were identified from the alignments using GATK v4.2.0.0 MarkDuplicates and excluded from downstream variant calling. The removal of duplicates and other aberrant reads by GATK’s default filters yielded a mean realized coverage of 28X (averaged across sites and across individuals).

Genotype likelihoods were computed for each individual using GATK v4.2.0.0 HaplotypeCaller with the Reference Confidence Model set to “GVCF” mode (Poplin et al. 2017). To accommodate between-population differences in allele frequencies and ensure well-calibrated measures of variant confidence, individual call sets were combined according to sampling location with GATK v4.2.0.0 CombineGVCFs and joint variant calling was performed separately for the Bumpkin (n=37), Peddocks (n=21), and World’s End (n=19) cohorts using GATK v4.2.0.0 GenotypeGVCFs (Poplin et al. 2017). Prior to filtering, our variant calling approach yielded a total of 51,857,593 SNPs in the Bumpkin cohort, 51,225,571 SNPs in the Peddocks cohort, and 74,861,232 SNPs in the World’s End cohort.

Due to a lack of available high-confidence call sets for training more sophisticated variant filtering approaches like Variant Quality Score Recalibration (VQSR), we elected to filter our variant calls based on a combination of site-level and individual-level annotations. GATK v4.2.0.0 VariantFiltration was used to remove variants based on the Best Practices recommendations for hard filtering (“FisherStrand” > 60.0, “StrandOddsRatio” > 3.0, “RMSMappingQuality” < 40.0, “MappingQualityRankSumTest” <-12.5, and “ReadPosRankSumTest” <-8.0; Van der Auwera and O’Connor 2020) with some minor modifications (“QualByDepth” < 5.0). Given that our downstream inferences about population structure and demographic history rely on accurate measures of sample allele frequencies, we additionally removed sites for which any individual’s read depth fell below 10X to ensure the accuracy of individual genotype calls.

Since low-complexity and repetitive regions of the genome can pose issues for read mapping and variant calling (Pfeifer 2017), we restricted our population genetic analyses to variants falling outside of annotated repetitive elements. We used the union of the RepeatMasker (hub_2100979_repeatMasker*; https://repeatmasker.org), TandemRepeatFinder (hub_2100979_simpleRepeat; Benson 1999), and WindowMasker (hub_2100979_windowMasker; Morgulis et al. 2006) annotation tracks of the UCSC Genome Browser (http://genome.ucsc.edu; Karolchik et al 2004) to exclude variants falling within these regions. Copy number variation that is poorly resolved in the reference assembly could instead manifest as regions of artificially inflated coverage (Li 2014; Pfeifer 2017). To mitigate the impact of unannotated repeats on the accuracy of variant calls, we additionally imposed a maximum coverage filter by removing sites for which any individual’s read depth exceeded 50X. Together, these quality-based and accessibility-based filtering criteria yielded a total biallelic SNP count of 7,964,169 in the Bumpkin Island cohort, 9,573,228 in the Peddocks Island cohort, and 12,990,178 in the mainland World’s End cohort.

#### Rescuing invariant sites

When there is sufficient allele frequency differentiation between populations, the joint calling approach described above will yield incongruent call sets, as variants that are private to one cohort will not be observed among the variant calls produced by other cohorts. This creates ambiguity about the status of missing variants, which could either represent monomorphic reference sites or sites for which there is insufficient information to obtain high-confidence genotypes. To ensure that our joint allele frequencies were properly calibrated for downstream analyses, we reinspected the genotypes at these missing sites.

Briefly, we used the bcftools v1.8 (Danecek et al. 2021) “isec” function to extract the locations of private variants within each pairwise comparison of call sets. We then produced VCF records for these missing sites within each cohort using the “-all-sites” mode of GATK GenotypeGVCFs with the “-stand-call-conf” threshold set to “0.0” to emit monomorphic reference calls. GATK v4.2.0.0 SelectVariants was used to restrict these “rescued” calls to invariant sites and the same site-level and individual-level filtering thresholds described above (excluding “QualByDepth”) were applied to obtain a collection of high-confidence, monomorphic calls. These rescued invariant calls were then merged with their respective variant calls, yielding a collection of data sets that represent the high-confidence union of variant sites observed across the three cohorts.

## Code availability

The pipeline used to process raw sequencing reads, perform the alignment, and conduct the variant calling and variant quality filtering is available on GitHub (https://github.com/PayseurLabUWMadison/boha_demography/tree/main/variant_calling).

### Exclusion of close relatives

To mitigate the potential biases that closely related individuals can introduce into population genetic analyses (Wang 2017), we estimated kinship coefficients between all pairs of individuals within each population sample using KING v2.2.8 (Manichaikul et al. 2010). Prior to kinship analysis, each variant call set was pruned for LD by removing SNPs with pairwise r^2^ > 0.1 in 50 bp sliding windows using the “--indep-pairwise 50 10 0.1” option in plink v1.90 (Chang et al. 2015). This LD-pruning yielded a total of 1,987,614 SNPs in the Bumpkin Island cohort, 1,253,128 in the Peddocks Island cohort, and 2,240,885 in the mainland World’s End cohort remaining for kinship analysis. The KING-Robust algorithm was used to estimate pairwise kinship coefficients from the LD-pruned autosomal variant calls and relationships were inferred from these estimates using the bounds provided by KING (Manichaikul et al. 2010). We used the “relatednessFilter” function of the R package plinkQC v0.3.4 to identify the maximum set of unrelated individuals (defined here as pairs ≥ 3^rd^ degree relatives) within each sample.

### Analysis of population structure

We analyzed population structure both between and within geographic locations by clustering individuals according to genetic similarity and by inferring individual ancestry proportions. Since both analyses assume that different SNPs experience free recombination, we conducted the same LD pruning described above within the Bumpkin, Peddocks, and World’s End cohorts. After excluding close relatives, this LD-pruning left a total of 92,099 SNPs in the Bumpkin cohort, 112,369 in the Peddocks cohort, and 213,660 in the World’s End cohort remaining for population structure analysis. PCA was performed on the combined, LD-pruned, autosomal call sets using the “--pca” option in plink v1.90 (Chang et al. 2015). Ancestry proportions were inferred either for each population sample separately or for the combined cohort with ADMIXTURE v1.3.0 (Alexander et al. 2009) using the same collection of SNPs. Five-fold cross-validation was performed for a range of ancestral components (k=1 to k=5) using the “--cv” option and individual ancestry proportions were inferred for the k value that yielded the lowest cross-validation error.

### Demographic inference

#### Model fitting and parameter estimation

We estimated demographic parameters using the maximum likelihood framework implemented in *Moments* v1.1.15 (Jouganous et al. 2017), which employs a recursion approach to compute the expected sample site frequency spectrum under a given model. To simplify the space of possible models and to ensure sufficient information in each shared allele frequency bin, we restricted our analysis to two-population models of divergence from a common ancestor. Tested models included either discrete or continuous changes in N_e_ for both island and mainland populations (Supplementary figure S6A). On top of these size changes, we fit two additional classes of models that allowed for either symmetric or asymmetric migration between sampled populations (Supplementary figures S6B and S6C, respectively). Given our limited sampling of the archipelago, we considered the possibility that observed variation might also reflect contributions made by unsampled populations inhabiting the many islands and peninsulas that comprise the Boston Harbor. To address this possibility, we attempted to fit a third class of migration models that allowed for symmetric migration between sampled populations and an unsampled “ghost” population (Supplementary figure S6D). To reduce the number of parameters in the ghost migration models, we held the ghost population’s N_e_ constant at the ancestral N_e_, assumed that it diverged from the ancestral population at the same time as the sampled populations, and assumed symmetric migration rates between the ghost population and focal population. For these three-population ghost migration models, we used the “marginalize” function to obtain the expected sample frequency spectrum for the two sampled populations.

To circumvent issues of ancestral state misidentification, we fit these models to the folded jSFS constructed from putatively neutral SNPs for each pair of populations. To increase the accuracy of estimates of sample allele frequencies, sites containing missing genotypes for one or more individuals were excluded from analysis. To mitigate the impact of direct and linked selection on observed allele frequencies, we masked annotated genes using the NCBI RefSeq annotations of the *P. leucopus* (UCI_PerLeu_2.1; GenBank accession: GCA_004664715.2; Long et al. 2019) reference assembly. Starting with our high-confidence variant callsets described above, we used the NCBI RefSeq track (hub_2100979_ncbiRefSeq; Pruitt et al. 2014) of the UCSC Genome Browser (http://genome.ucsc.edu; Karolchik et al 2004) to exclude SNPs falling within 100 bp of predicted and curated genes. Given the potential for the effects of linked selection to extend farther distances from these functional regions, we tested the impact of more stringent distance-based gene masking on the observed jSFS for each pair of populations (Supplementary figures S7). We found that extending the masked distance beyond 100 bp upstream and downstream of gene annotations to distances of 5 kbp, 50 kbp, and even 100 kbp had little impact on the genome-wide jSFS constructed from the remaining SNPs (Supplementary figures S7). Given this finding, we limited our masking to 100 bp surrounding annotated genes to retain the maximal number of SNPs for inference. This approach left a total of 4,219,108 Bumpkin SNPs, 5,063,432 Peddocks SNPs, and 6,707,809 World’s End SNPs remaining for inference.

For each of the tested models, we used the BFGS optimization algorithm implemented in *Moments* to find the parameter values that maximized the likelihood of the observed jSFS. We permuted starting parameter values across 10,000 independent searches and retained the top three parameter combinations that yielded the highest likelihood for each model. Following model fitting, we employed a hierarchical approach to selecting between contrasting demographic models. First, we evaluated parameter convergence and eliminated models for which the coefficient of variation for any parameter exceeded 0.2 across the top three highest likelihood estimates. Next, we compared likelihoods between models, either directly for models with equivalent numbers of parameters, or through adjusted likelihood ratio tests (Coffman et al. 2016) for models with nested parameters. Finally, for remaining models, we inspected their fit to both the jSFS and the marginal SFS for each component population and proceeded with the simplest model that yielded the best fit to these frequency spectrum partitions.

For selected models, we calculated the ancestral effective population size as N_Anc_ = θ/(4*µ*L) where we assumed a mutation rate, µ, of 4e-9 per-site, per-generation and used an effective sequence length, L, that represents the total length of sequence from which we could have called variants. To obtain an approximate L of 210,600,000 bp, we multiplied the total length of the autosomal portion of the *P. leucopus* reference assembly (2,336,931,804 bp) by the average percent of called SNPs that survived our quality-based, accessibility-based, and neutrality-based filtering (9.01%). Population size and time parameters were scaled by N_Anc_ and 2*N_Anc_ to obtain these parameter estimates in units of individuals and generations, respectively. LD among SNPs used to construct the jSFS violates the independence assumption of the Poisson Random Field framework used to compute model likelihoods (Sawyer and Hartl 1992) and precludes the use of standard techniques to estimate the uncertainty of model parameters. As an alternative to the computationally intensive approach of conventional bootstrapping, we used the Godambe Information Matrix methods described in Coffman et al. (2016) to estimate parameter uncertainties in the presence of this non-independence. Since certain population size and time parameters are functions of multiple estimated quantities, we used uncertainty propagation techniques (Ku 1966) to obtain the standard deviations of these parameter estimates.

#### Predictive simulations

We evaluated the fit of the inferred demographic histories to other patterns of neutral variation by conducting predictive simulations with *msprime* v1.2.0 (Baumdicker et al. 2022). For each population pair, we simulated genomic data sets using the maximum likelihood parameter estimates obtained for the best-fit demographic model. These genomic data sets were constructed from 1,000 independent simulations of a 1 Mbp genomic element using a uniform recombination and mutation rate of 4e-9. Simulated sample sizes were set to match those of the empirical data. For both the simulated and empirical neutral data sets, we computed nucleotide diversity π (Tajima 1983), Tajima’s *D* (Tajima 1989), and F_ST_ based on nucleotide diversity (Hudson, Slatkin, and Maddison 1992) in non-overlapping 5 kbp physical windows using scikit-allel v1.3.6 (https://github.com/cggh/scikit-allel). To compare the decay of LD between simulated and observed data, we used the “--r2” option in plink v1.90 (Chang et al. 2015) to measure pairwise r^2^ between a random subset of SNPs sampled from simulated and empirical chromosomes. We restricted this analysis to SNPs separated by at most 1 Mbp by setting the “--ld-window” and “--ld-window-kb” arguments to 1,000,000 and 1,000, respectively. Singleton variants were excluded from LD calculations by setting the “--mac” argument to 1. The rigorous quality-based, accessibility-based, and neutrality-based filtering that we performed on the empirical data introduced substantial heterogeneity in the number of “observable” sites that comprise each 5 kbp physical window. To account for this, we computed per-site statistics like π based on the total number of observed sites using the “is_accessible” argument in scikit-allel calculations. For SNP-based statistics like Tajima’s *D* and F_ST_, we note that this heterogeneity could be a major source of between-window variability in summary statistic estimates from the empirical data.

## Code availability

The scripts used to conduct demographic inference with *Moments* and to perform the predictive simulations with *msprime* are available on GitHub (https://github.com/PayseurLabUWMadison/boha_demography/tree/main/demographic_infer ence).

### Singleton analysis

Inspection of variant positions along chromosomes revealed clustering of singleton variants by mouse and by genomic location in the island samples. To explore the capacity of migration to generate this pattern, we used chromosome-scale simulations conducted with *msprime* v1.2.0 (Baumdicker et al. 2022). Given that best-fit demographic models based on the jSFS did not include migration, we augmented our best-fit demographic histories with gene flow from an unsampled “ghost” population (Supplementary figure S8). Our aim was to capture a range of scenarios that might be relevant to migration that occurs locally within the archipelago. To this end, we varied both the strength of migration between the focal populations and ghost population (spanning rates of 1e-3, 1e-4, and 1e-5) as well as the duration of time over which migration occurs, expressed as the proportion of the total split time estimated for each pair of sampled populations (either 0.1%, 1%, 10%, or 100% of the inferred split time) (Supplementary figure S8). To reduce the number of tested migration models, we made several simplifying assumptions. First, we assumed that the divergence of the ghost population from the ancestral population coincided with the divergence of the sampled populations, and we held the ghost N_e_ constant at the ancestral N_e_. Second, we assumed that both sampled populations experienced the same amount of migration from the same ghost population. Third, we assumed that migration between the sampled and ghost population occurred in a continuous and symmetric fashion into the present. For each migration regime, we simulated 25 independent 100 Mbp genomic elements using the parameter estimates from the best-fit demographic model for each pair of populations. For each simulation, we assumed a uniform recombination and mutation rate of 4e-9. Simulated sample sizes were set to match those of the empirical data. For comparison, we generated additional chromosome-scale data sets in the absence of migration.

We used the chi-squared test statistic to quantify the deviations of individual singleton contributions from the null hypothesis that singletons are contributed equally among individuals. For each of the simulated and empirical chromosomes, we used R v4.2.1 (R Core Team 2022) to compute the chi-squared test statistic from the observed and expected individual singleton counts. To evaluate the impact of migration on the jSFS between island and mainland populations, we used scikit-allel v1.3.6 (https://github.com/cggh/scikit-allel) to compute joint allele frequencies from datasets simulated with and without ghost migration.

To examine the combined effect of population size history and migration on the genealogies underlying the singleton pattern, we revisited the simulations of Bumpkin Island chromosomes conducted using the best-fit parameter estimates from the Bumpkin–World’s End demographic history (Figure 3) with migration parameters from the Recent/Strong regime featured in Figure 6B. We prioritized this model because it involves the most extreme reduction in N_e_ for the island population (Figure 3) and provided the closest qualitative fit to the observed singleton tracts (Figures 5A and 6B). We used tskit v0.5.8 (Kelleher et al. 2018) to inspect the properties of marginal trees spanning a single 100 Mbp Bumpkin Island chromosome. We used the “migrations” function to record the breakpoints of migrant ancestry along the chromosome. Then, we used the “trees” function to iterate over all of the marginal trees in the tree sequence. For each of the 1,010,931 trees, we compared the tree boundaries to the recorded breakpoints of migrant ancestry to determine if the tree overlapped one or more migrant haplotypes (classified as “with migration”) (Figure 7A). Trees were classified as “no migration” if they did not span migrant tracts (Figure 7A). We computed the branch length quantities illustrated in Figure 7B for each tree. We also recorded the TMRCA of each tree along with the population in which the common ancestor originated.

## Code availability

The scripts used to perform the migration simulations with *msprime* and analyze the simulation output with scikit-allel and tskit are available on GitHub (https://github.com/PayseurLabUWMadison/boha_demography/tree/main/migration_analysis)

### Estimating contemporary effective population sizes

We estimated contemporary effective population sizes in the Boston Harbor using two LD-based approaches, *currentNe* (Santiago et al. 2024) and GONE (Santiago et al. 2020). Observed levels of LD can be influenced by the structure of relatedness in a sample, and there is evidence that excluding relatives may artificially inflate estimates of contemporary effective population size obtained by these methods (Waples and Anderson 2017; Santiago et al. 2024). Because we cannot know whether the abundance of close relatives we identified (Supplementary figure S1) reflects random sampling or an artifact of trapping procedure, we chose to conduct these analyses on both the complete sample and on the subset of unrelated individuals from each population (obtained by excluding relationships ≤ 2^nd^ degree). For all analyses of the Bumpkin Island population, we additionally excluded individual B19 due to its genetic divergence from the rest of the sample (Supplementary figures S2, S4, and S5). We used the same putatively neutral, high-confidence, autosomal SNP callset used for demographic inference with *Moments* as the input to these analyses (see sections ***Sequence data processing and variant calling*** and ***Demographic inference***).

For each dataset, we obtained point estimates of N_e_ reflecting the few most recent generations with *currentNe* (Santiago et al. 2024). For computational tractability, we used the “-s” argument to restrict pairwise measurements of LD to 50,000 randomly sampled SNPs. N_e_ estimates were based only on pairs of SNPs located on different chromosomes. To account for the impact of close relatives, we used the “-k” option to jointly estimate N_e_ and k, the average number of full siblings each individual has in the sample (Santiago et al. 2024). Consistent with our KING analyses (Supplementary figure S1), this parameter yielded estimates of 0.53, 0.43, and 0.19 for the complete Bumpkin Island, Peddocks Island, and mainland World’s End samples, respectively. As expected, we obtained k values of 0 for each sample of unrelated individuals. 90% confidence intervals for each point estimate were obtained using the artificial neural network (ANN) implemented in the *currentNe* software (Santiago et al. 2024).

To reconstruct the recent trajectory of N_e_ in each population, we analyzed each dataset with GONE (Santiago et al. 2020). We used plink v1.90 (Chang et al. 2015) to convert input SNP callsets from VCF to plink text files (.ped and.map) with the “--recode” option. Singleton variants and sites with missing genotype calls were excluded from each analysis by setting the “--mac” argument to 1 and the “--geno” argument to 0. For all GONE analyses, we used the default settings for unphased genotypes and specified a uniform recombination rate of 0.4 cM/Mb to match the rate assumed for predictive simulations (see ***Demographic inference***). For each time interval, we report the geometric mean N_e_ estimate obtained across 40 replicates. In contrast to *currentNe*, GONE does not explicitly account for the presence of close relatives in the dataset (Santiago et al. 2024). Despite this, the impact of close relatives is expected to be negligible, since they exert greater influence on LD between unlinked markers than between the closely linked markers used by GONE to estimate the recent N_e_ trajectory (Santiago et al. 2024).

## Acknowledgements

We thank members of the Payseur lab for their helpful input on this work. We thank the National Park Service for assistance with boat transportation, mouse capture, and sample collection. This research was funded by National Institutes of Health (NIH) grants R01GM100426 and R35GM139412 (to B.A.P.). E.K.H. was partially supported by the NIH Graduate Training Grant in Genetics at the University of Wisconsin-Madison (T32GM007133). Computational analyses were conducted using resources provided by the University of Wisconsin-Madison’s Center for High Throughput Computing.

## Data Accessibility

The code used to conduct all analyses is provided on GitHub (https://github.com/PayseurLabUWMadison/boha_demography). Upon submission, raw sequencing reads will be deposited in the SRA and a static snapshot of the code will be archived on Dryad.

## Author Contributions

E.K.H., B.A.P., and L.E.N. designed the study. L.E.N. acquired the samples used for the study.

E.K.H. conducted the research with supervision from B.A.P. E.K.H. and B.A.P. wrote the manuscript with input from L.E.N.

