## Supplementary for "Population history across timescales in an urban archipelago"

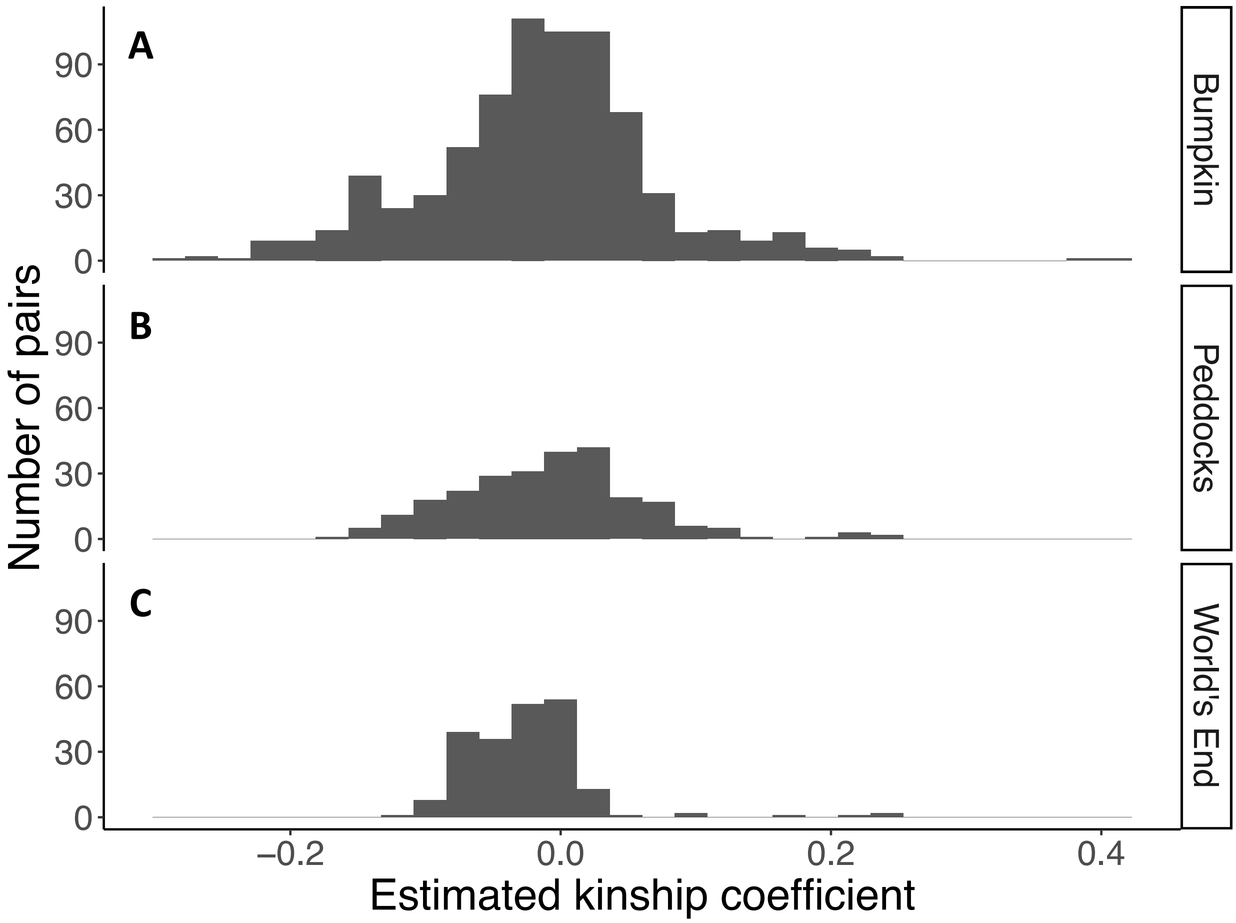


**Figure S1: High incidence of close relatives in the island samples.** Pairwise kinship coefficients estimated by KING for Bumpkin Island (A), Peddocks Island (B), and mainland World's End (C) individuals. X-axis gives the kinship coefficient estimates produced by KING and Y-axis gives the number of pairs observed for a given kinship coefficient bin.

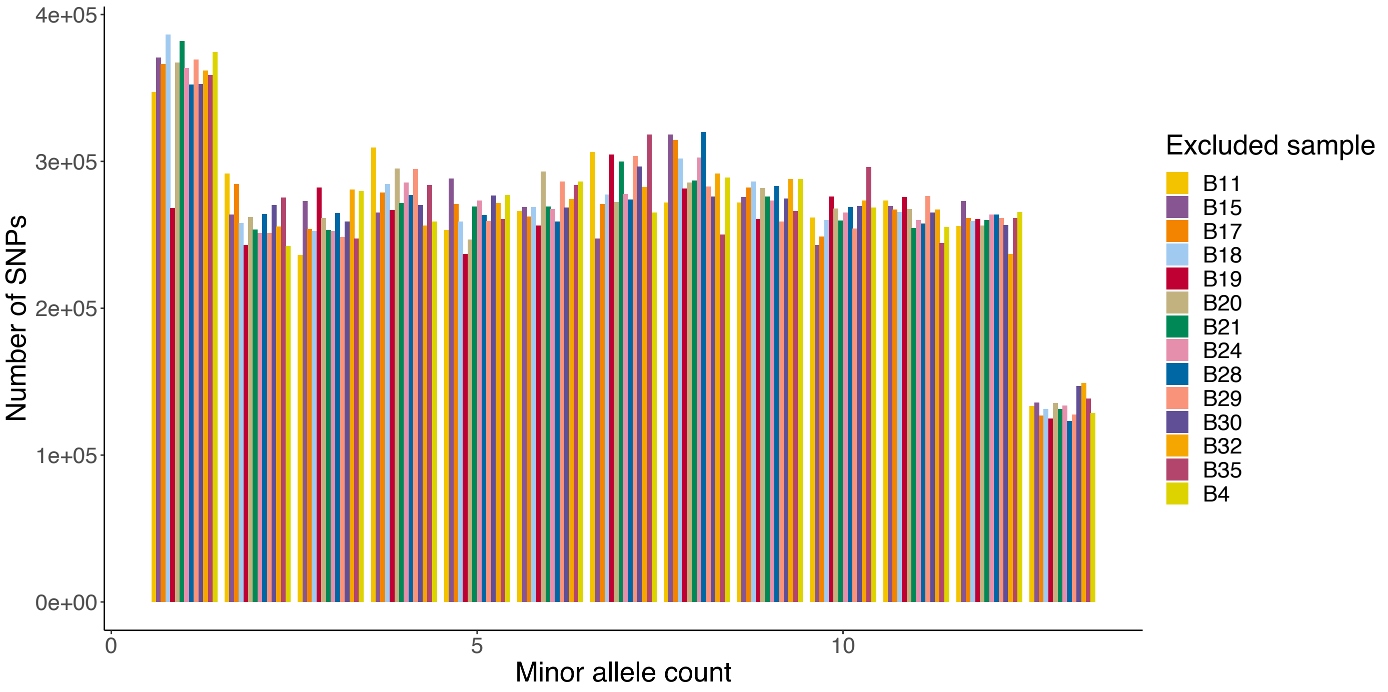


**Figure S2: A single Bumpkin individual contributes a disproportionate number of singletons.** Leave-one-out resampling of the folded site frequency spectrum (SFS) revealed that the exclusion of individual B19 (red) caused a large reduction in the number of observed singleton variants (i.e., those with a minor allele count of 1). The y-axis gives the number of observed SNPs for each minor allele count bin on the x-axis. Re-sampled frequency spectra are colored according to the individual that was excluded.

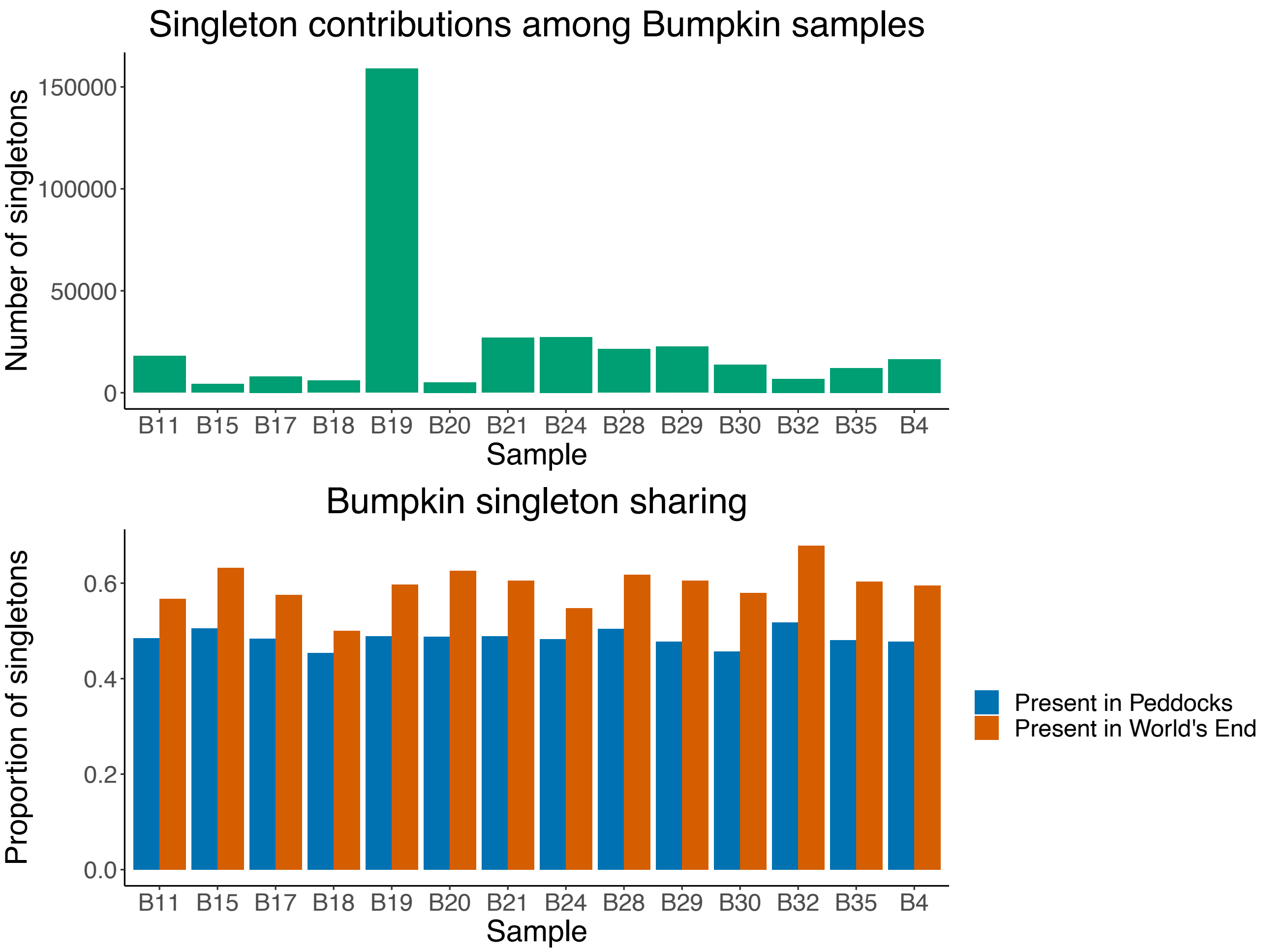


**Figure S3: Singletons found in divergent Bumpkin individual are shared with the Peddocks and mainland World's End cohort.** (Top) number of singletons (y-axis) contributed by each Bumpkin Island individual (x-axis) to the folded site frequency spectrum (SFS). (Bottom) Proportion of each individual's contributed singletons (y-axis) that are found in either the Peddocks Island (blue) or mainland World's End (orange) cohort. Despite contributing a disproportionate number of singletons to the sample, a similar proportion of individual B19's singletons are shared with the other cohorts, suggesting that genotyping error is an unlikely source for these excess singletons.

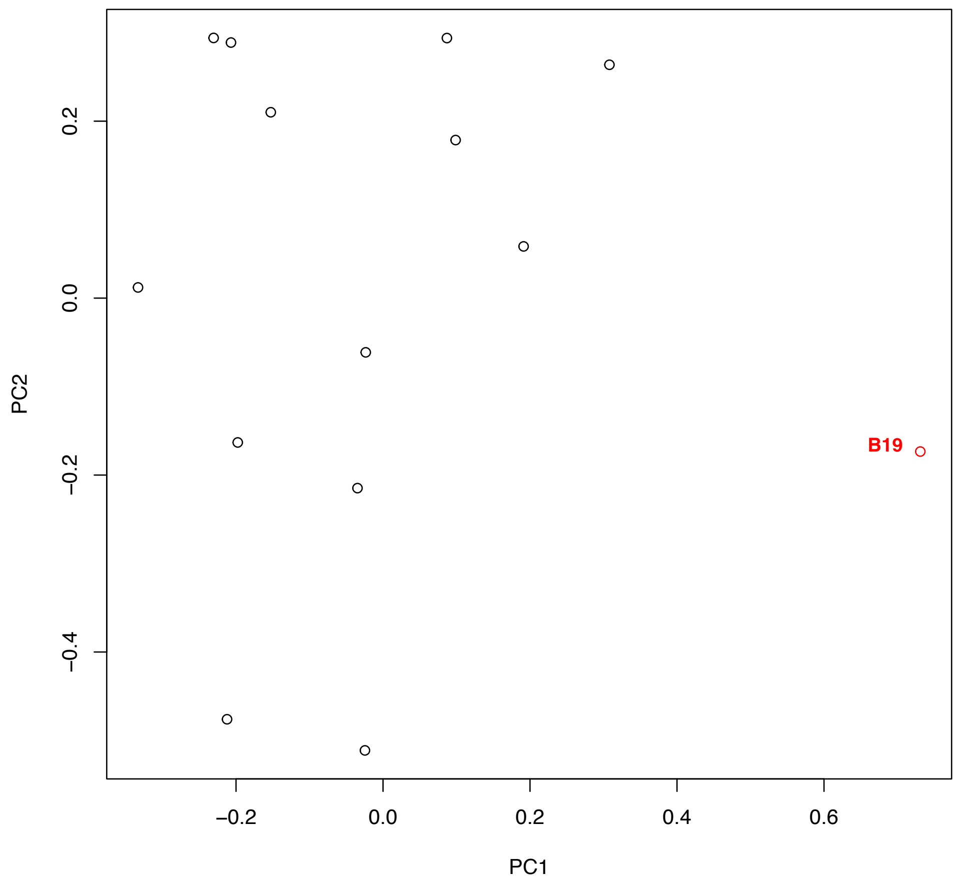


**Figure S4: Individual B19 exhibits departures from the Bumpkin Island sample.** Principal component analysis (PCA) conducted using genome-wide, unlinked SNPs reveals divergence between individual B19 (red point) and other Bumpkin Island individuals (black points). Percentage of variance explained by PC1=13.02% and by PC2=10.60%.

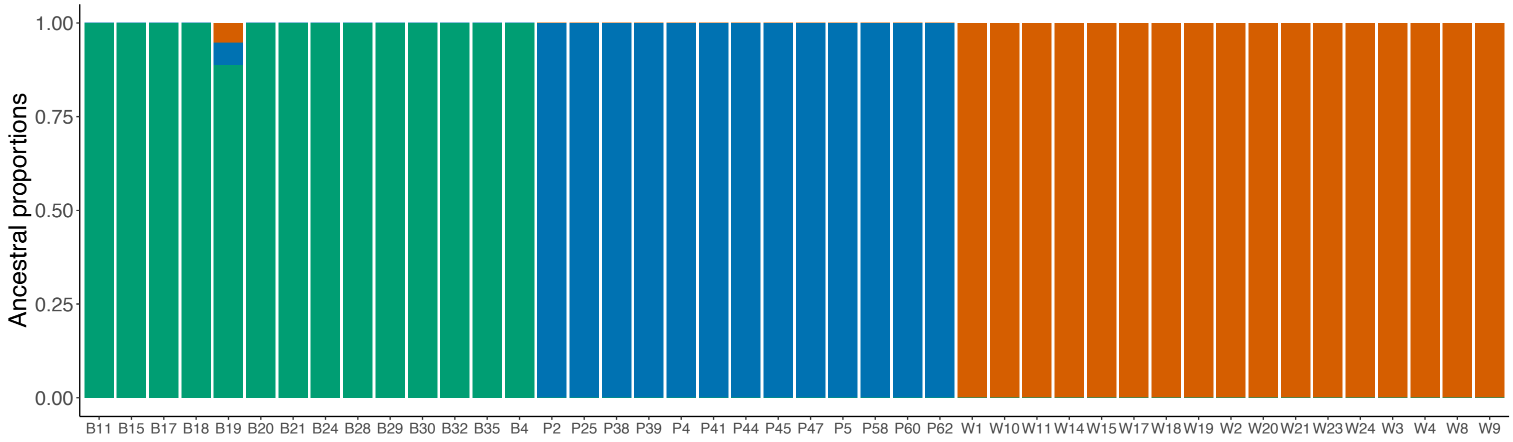


**Figure S5: Bumpkin Island individual B19 shares ancestry with other Boston Harbor locales.** Global ancestry proportions among unrelated individuals assuming k=3 ancestral components. Each individual is represented by a colored bar. The height of each colored segment in a bar gives the proportion of ancestry an individual derives from a given component. Ancestral component colors reflect population of origin: Bumpkin Island (green), Peddocks Island (blue), or mainland World's End (orange). Unlike other Bumpkin Island mice, individual B19 (fifth bar from left) derives ancestry from both the Peddocks Island and mainland World's End components.

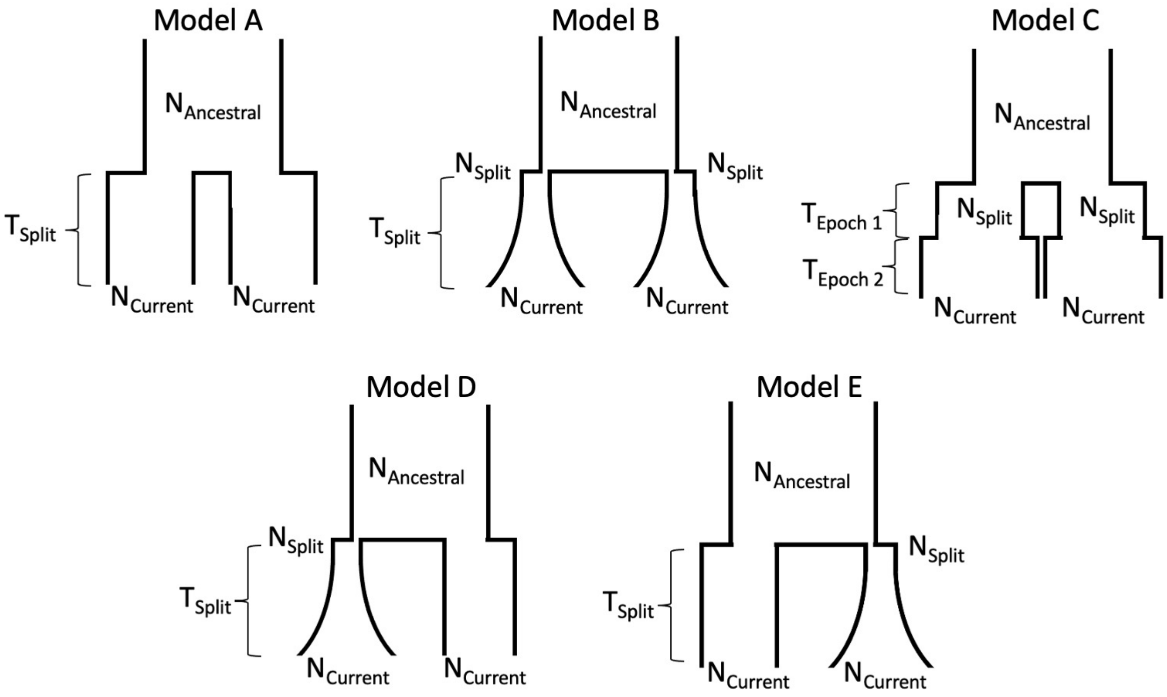


**Figure S6A: Two-population demographic models.** Models A-E correspond to simplified models of population splits that involve either discrete (Models A and C), continuous (Model B), or a mixture of discrete and continuous (Model D and E) changes in effective population size following the split. Continuous changes in effective population size were modeled as exponential growth (or decline). Discrete changes were modeled as instantaneous growth (or decline). The direction of size change was not constrained by the model. All estimated parameters are labeled for each of the tested models.

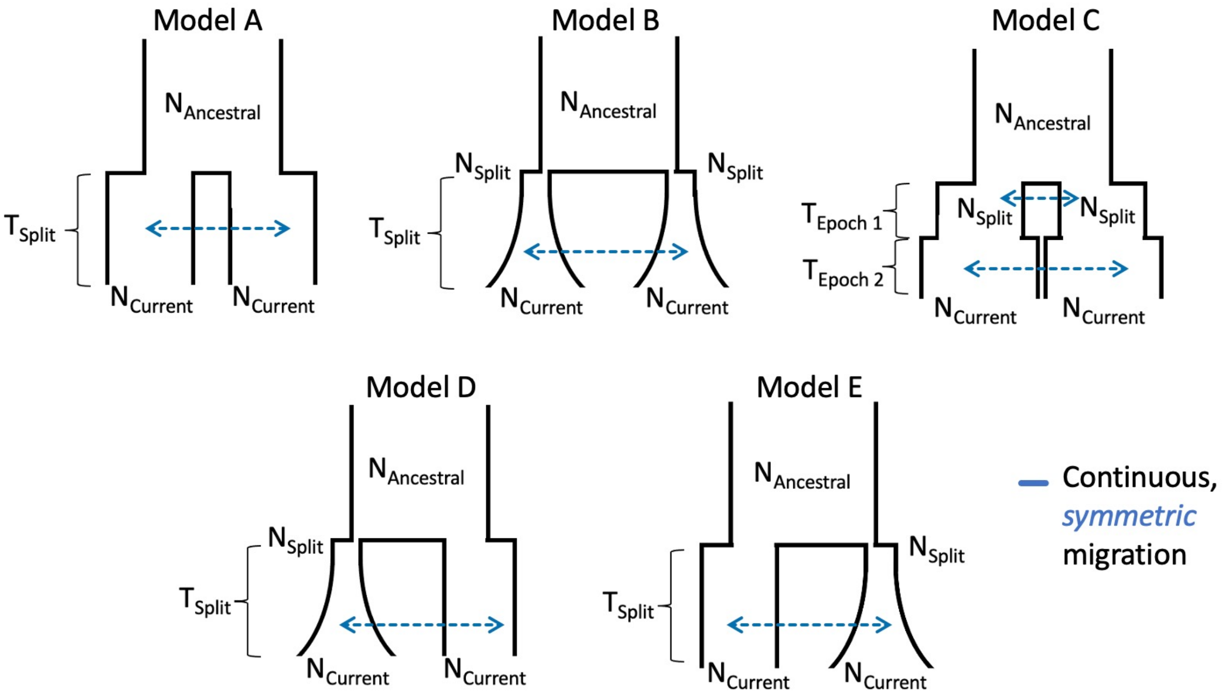


**Figure S6B: Two-population demographic models with symmetric migration.** Models A-E are specified as in Figure S6A with the addition of either one (Models A, B, D, and E) or two (Model C) migration rate parameters that model continuous, symmetric migration occurring between the two sampled populations after their divergence from a shared ancestor.

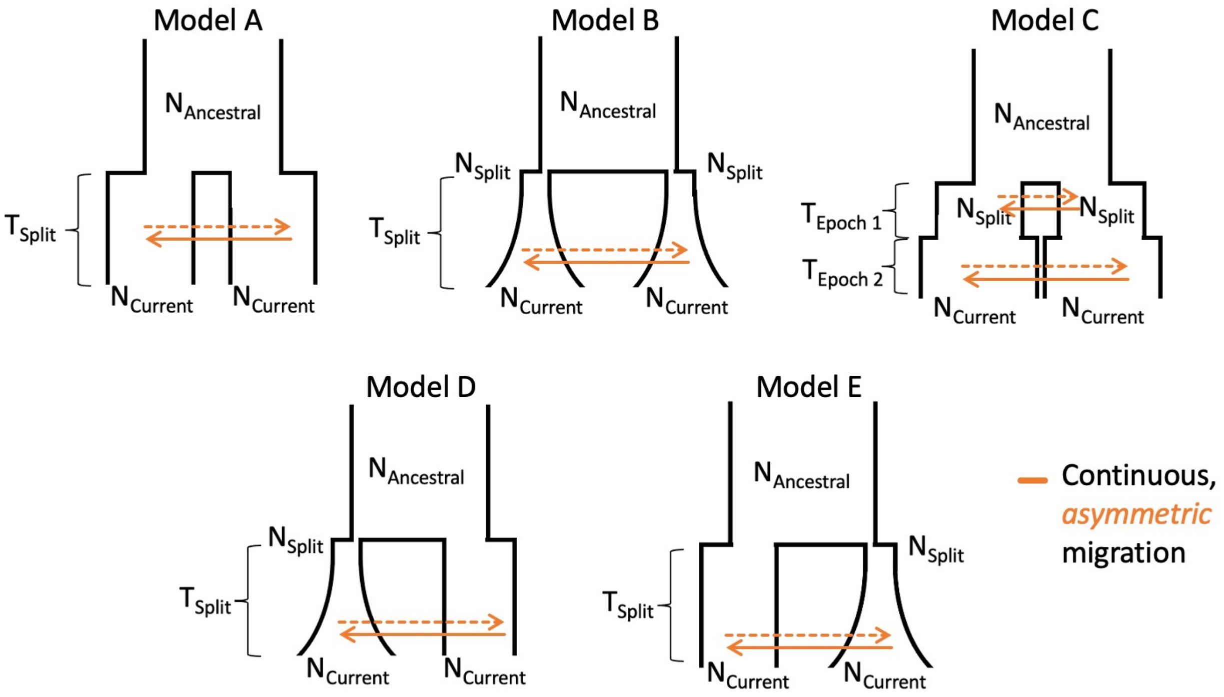


**Figure S6C: Two-population demographic models with asymmetric migration.** Models A-E are specified as in Figure S6A with the addition of either one (Models A, B, D, and E) or two (Model C) pairs of migration rate parameters that model continuous, asymmetric migration occurring between the two sampled populations after their divergence from a shared ancestor.

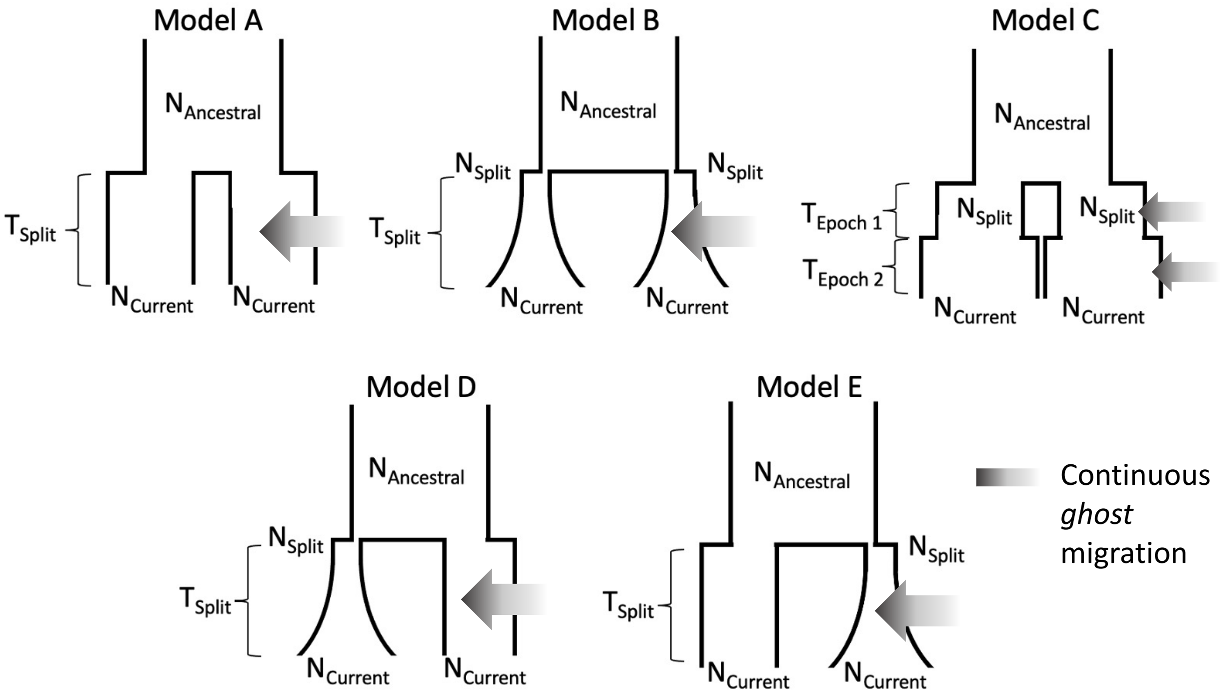


**Figure S6D: Two-population demographic models with ghost migration.** Models A-E are specified as in Figure S6A with the addition of either one (Models A, B, D, and E) or two (Model C) migration rate parameters that model continuous, symmetric migration between the specified focal population and an unsampled "ghost" population. The ghost population's size was held constant at $N_{Ancestral}$ and we assumed that it diverged from the ancestral population at the same time as the two sampled populations.

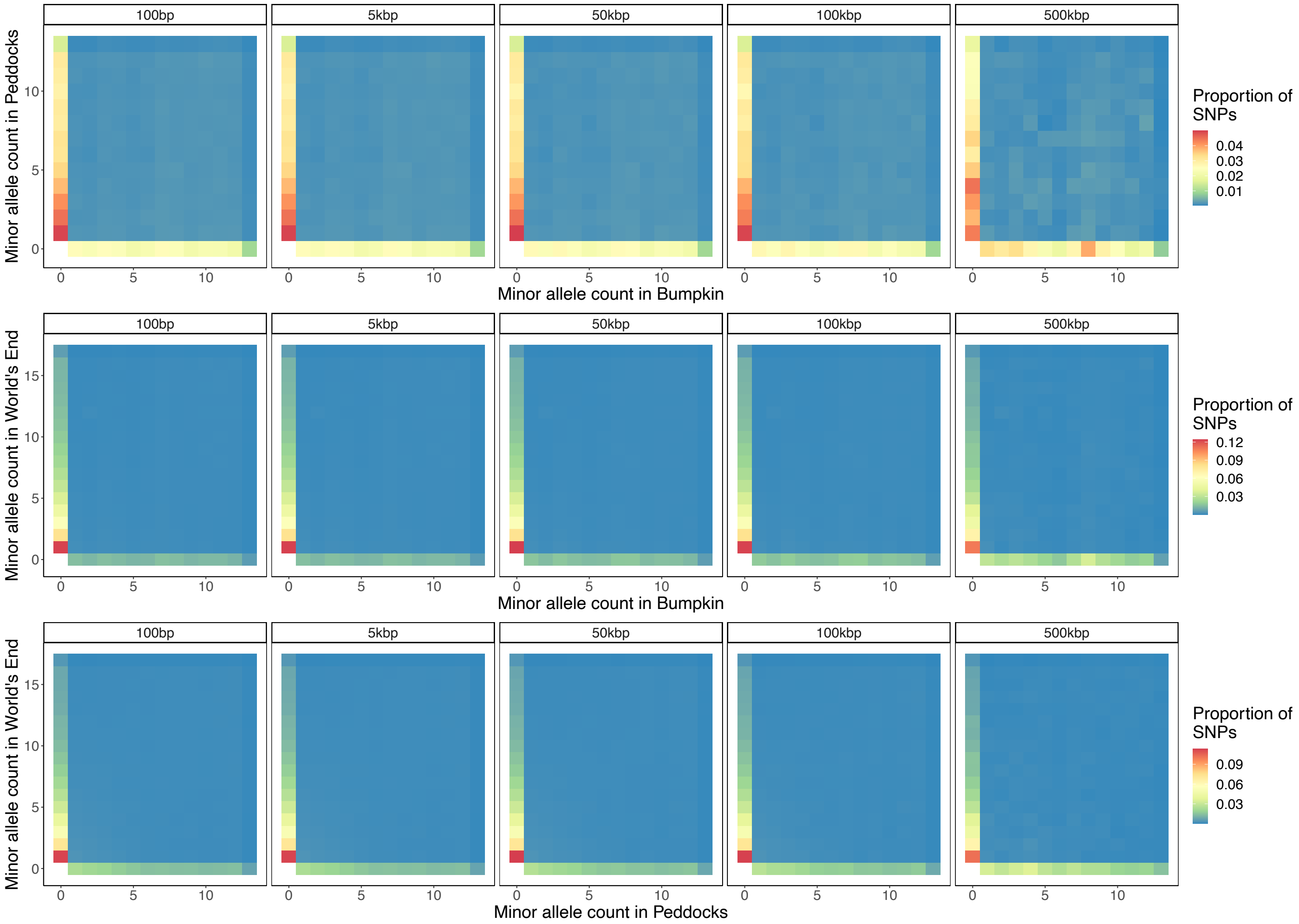


**Figure S7: Limited effect of distance-based gene masking on joint allele frequencies.** Joint site frequency spectra (jSFS) for the Peddocks Island-Bumpkin Island (top), mainland World's End-Bumpkin Island (middle), and mainland World's End-Peddocks Island (bottom) comparisons. Each jSFS was constructed by excluding single nucleotide polymorphisms (SNPs) falling within or near NCBI RefSeq gene annotations. Plots are facetted according to the distance from gene annotations at which SNPs were removed (left to right: 100 bp, 5 kbp, 50 kbp, 100 kbp, or 500 kbp). The y-axis gives the minor allele count of an observed SNP in population 1 and the x-axis gives the minor allele count of the same SNP in population 2. The color of each cell reflects the proportion of total SNPs observed in the given frequency bin.

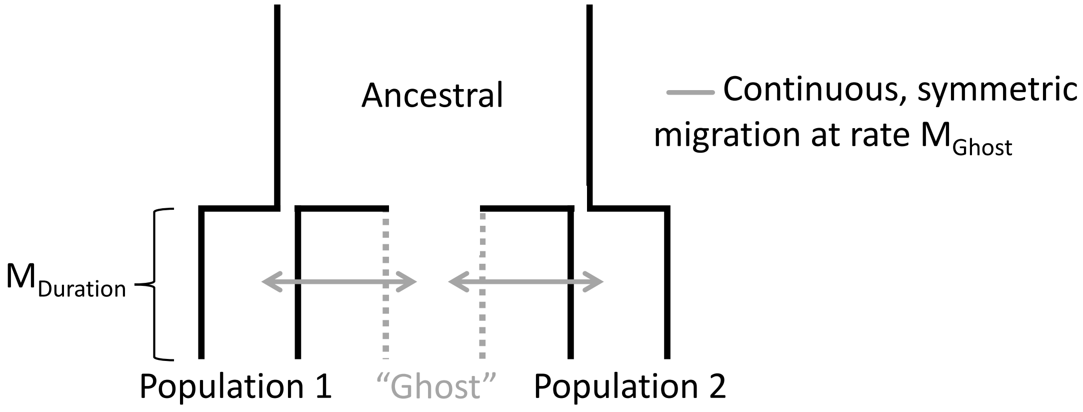


**Figure S8: Demographic model used for migration simulations.** We augmented the best-fit models of population divergence obtained from our demographic analyses with various scenarios of migration from an unsampled "ghost" population. Using the estimated $N_{Ancestral}$, $N_{Population\ 1}$, $N_{Population2}$, and $T_{Split}$ for each population pair as the foundation for our simulations, we tested a series of symmetric migration rates (1e-3, 1e-4, and 1e-5) between the ghost population and each sampled population. We also varied the total duration over which migration occurs (expressed as either 0.1%, 1%, 10%, or 100% of $T_{split}$). The ghost population's size was held constant at $N_{Ancestral}$, and we assumed that it diverged from the ancestral population at the same time as the two sampled populations. We also assumed the same rate of migration between the ghost population and each of the sampled populations.

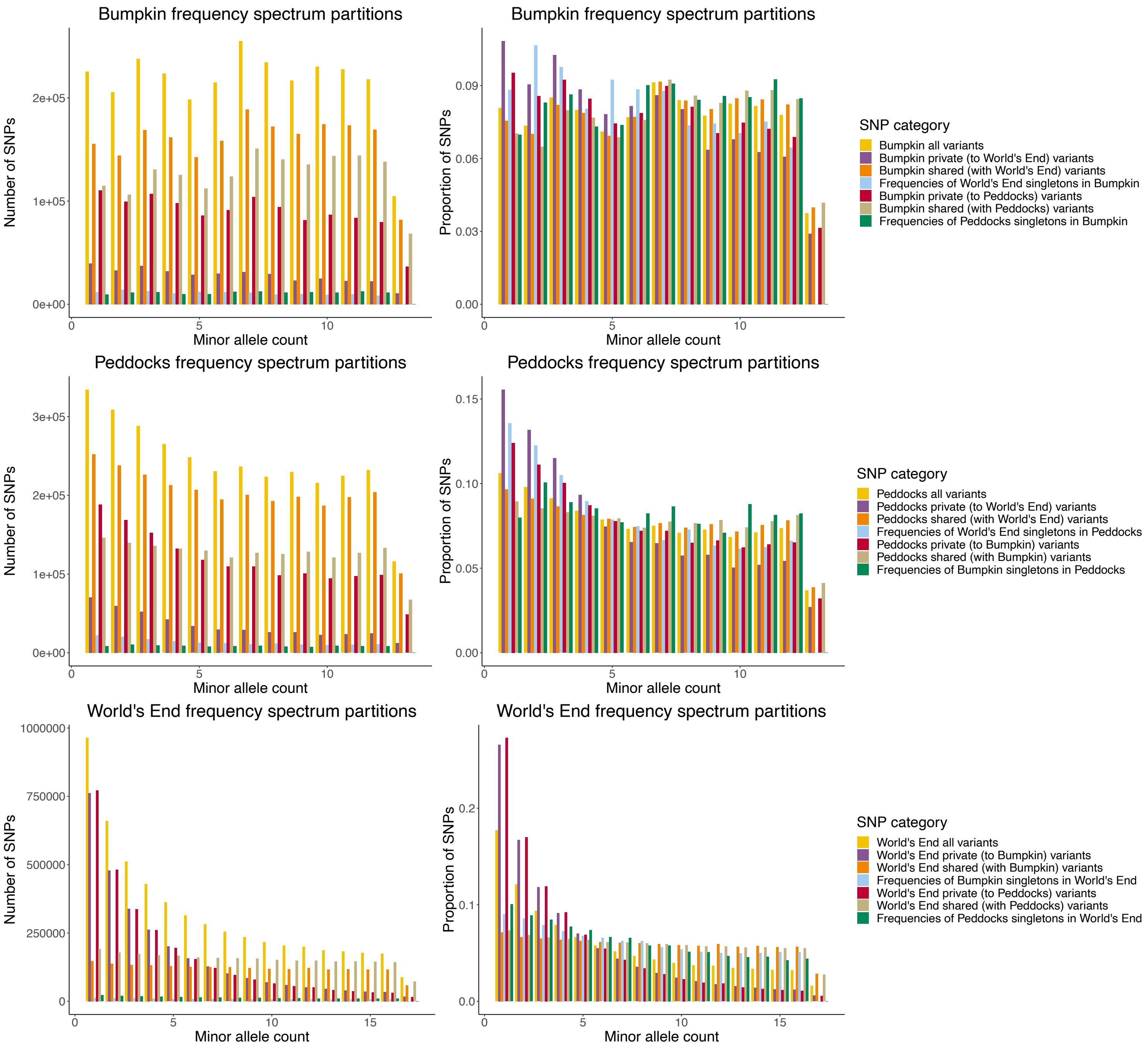


**Figure S9: Site frequency spectrum partitions.** Folded site frequency spectra of Bumpkin Island (top), Peddocks Island (middle), and mainland World's End (bottom) samples. Colored bars reflect the frequency spectrum of different partitions of shared (orange and beige), private (purple and red), and shared singleton (blue and green) variants in the focal population. For comparison to these partitions, the complete folded SFS of the focal population is plotted in yellow. (Left) SFS plot in terms of the total number of SNPs. (Right) SFS plot in terms of the proportion of SNPs in each frequency bin.

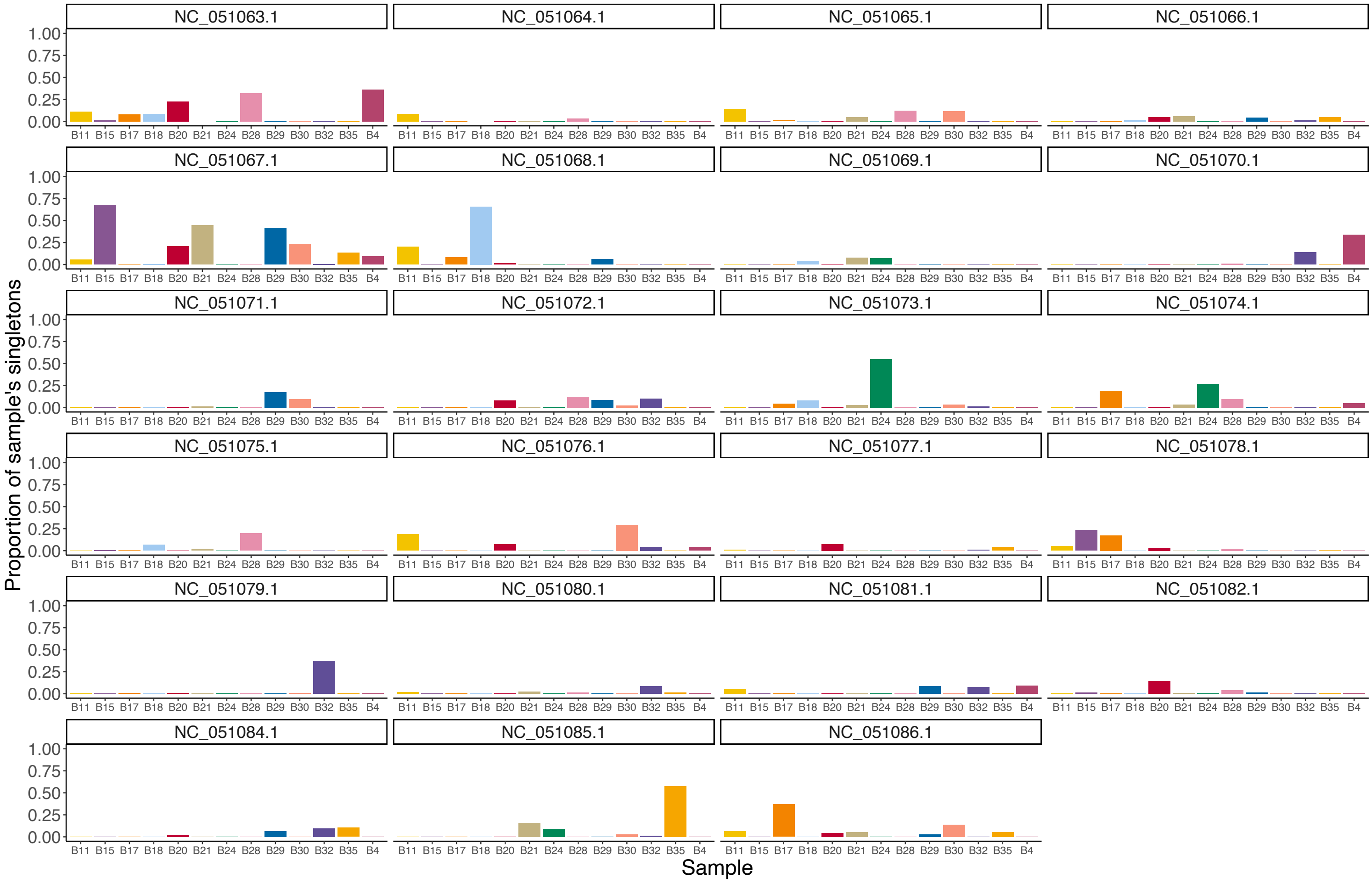


**Figure S10: Bumpkin Island individuals demonstrate high inter-chromosomal variance in singleton number.** Colored bars plot the proportion of each Bumpkin Island individual's total contributed singletons found on a given chromosome. Plots are faceted according to autosome using RefSeq accession IDs.

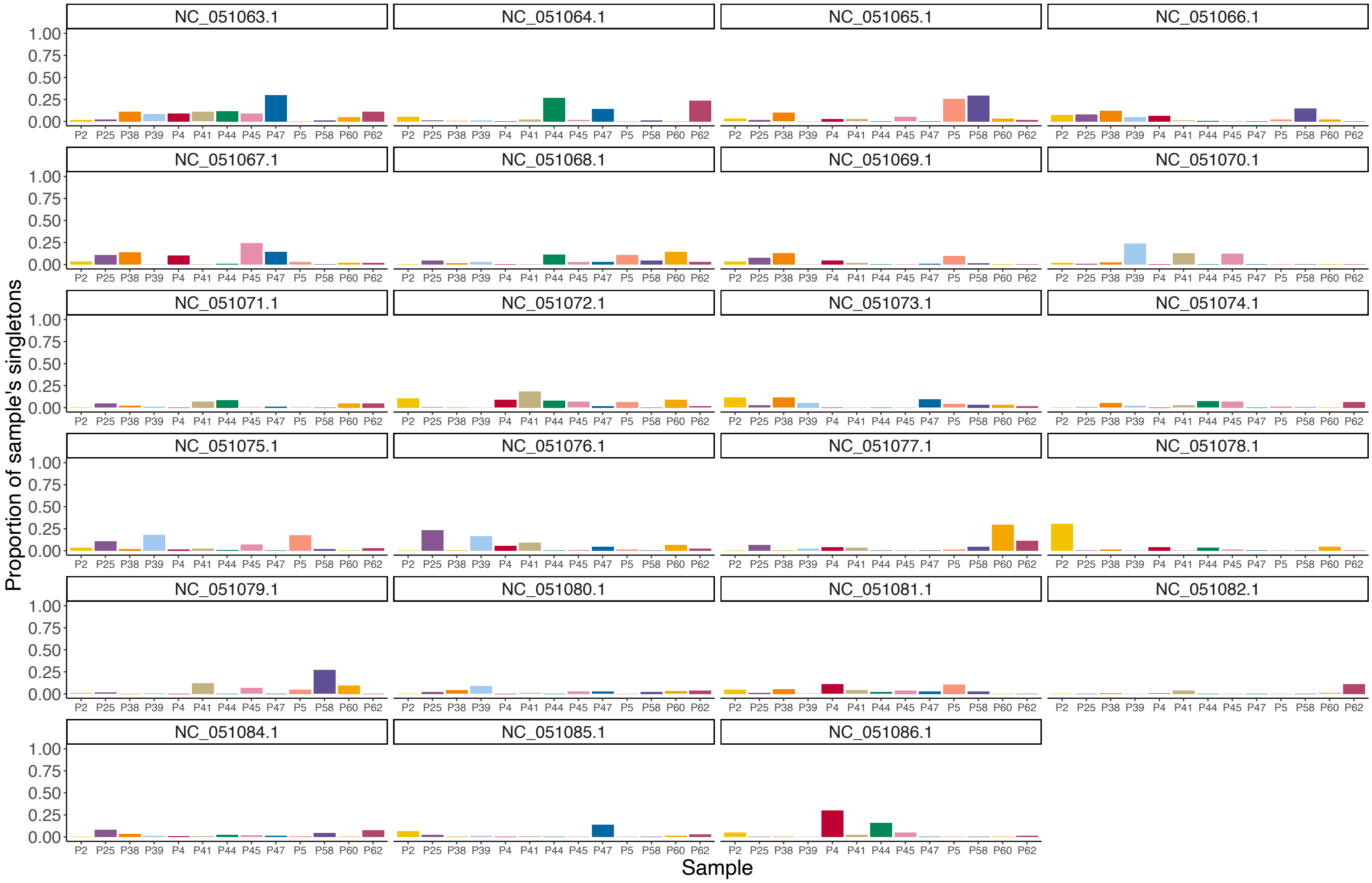


**Figure S11: Peddocks Island individuals demonstrate high inter-chromosomal variance in singleton number.** Colored bars plot the proportion of each Peddocks Island individual's total contributed singletons found on a given chromosome. Plots are faceted according to autosome using RefSeq accession IDs.

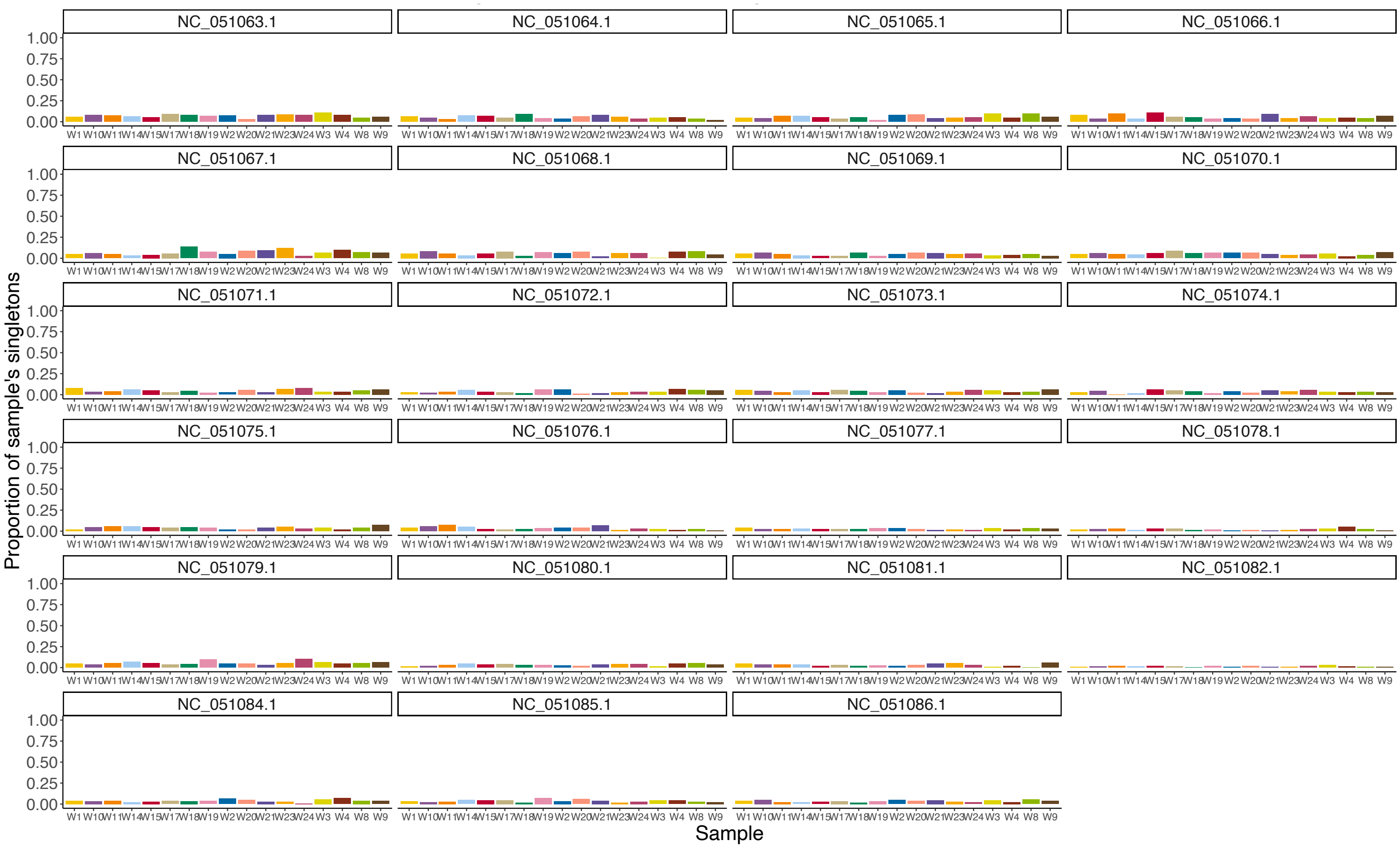


**Figure S12: Mainland World's End individuals demonstrate low inter-chromosomal variance in singleton number.** Colored bars plot the proportion of each mainland World's End individual's total contributed singletons found on a given chromosome. Plots are faceted according to autosome using RefSeq accession IDs.

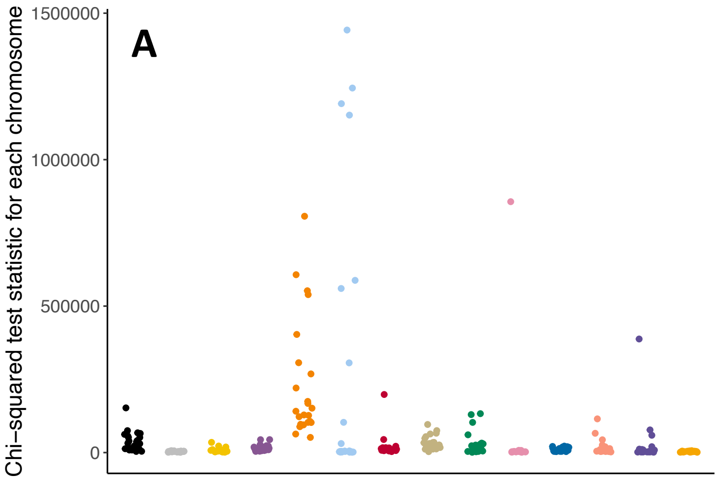


| Population | Params | Mean | Sd |
|---|---|---|---|
| Bumpkin | Empirical | 38305.425 | 33444.995 |
| Bumpkin | MigRate−0; MigTime−0 | 3562.222 | 1427.468 |
| Bumpkin | MigRate−1e−3; MigTime−1e−0 | 7795.111 | 7836.176 |
| Bumpkin | MigRate−1e−3; MigTime−1e−1 | 15749.416 | 10655.766 |
| Bumpkin | MigRate−1e−3; MigTime−1e−2 | 229906.456 | 202963.850 |
| Bumpkin | MigRate−1e−3; MigTime−1e−3 | 266719.556 | 472668.784 |
| Bumpkin | MigRate−1e−4; MigTime−1e−0 | 20299.003 | 38766.690 |
| Bumpkin | MigRate−1e−4; MigTime−1e−1 | 33280.389 | 22235.066 |
| Bumpkin | MigRate−1e−4; MigTime−1e−2 | 29096.547 | 37696.310 |
| Bumpkin | MigRate−1e−4; MigTime−1e−3 | 37760.204 | 170571.974 |
| Bumpkin | MigRate−1e−5; MigTime−1e−0 | 10115.373 | 6205.130 |
| Bumpkin | MigRate−1e−5; MigTime−1e−1 | 15967.825 | 25167.417 |
| Bumpkin | MigRate−1e−5; MigTime−1e−2 | 25310.322 | 77664.171 |
| Bumpkin | MigRate−1e−5; MigTime−1e−3 | 3409.065 | 1352.605 |

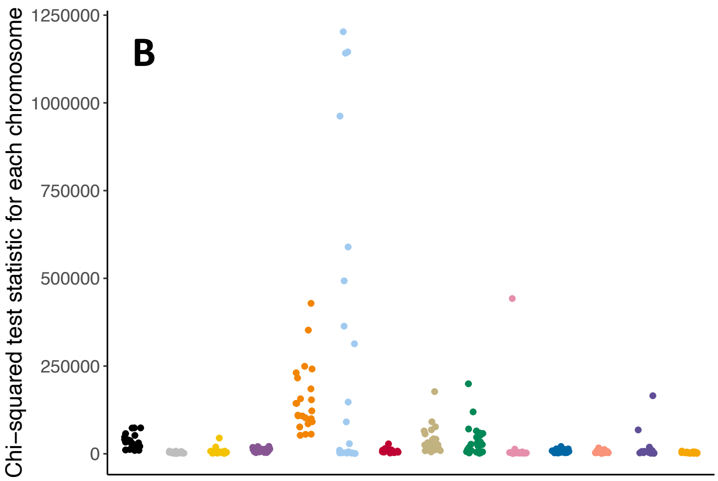


| Population | Params | Mean | Sd |
|---|---|---|---|
| Peddocks | Empirical | 34745.252 | 20402.148 |
| Peddocks | MigRate−0; MigTime−0 | 3506.979 | 1874.080 |
| Peddocks | MigRate−1e−3; MigTime−1e−0 | 6326.696 | 8755.343 |
| Peddocks | MigRate−1e−3; MigTime−1e−1 | 10459.236 | 5243.267 |
| Peddocks | MigRate−1e−3; MigTime−1e−2 | 152731.917 | 93922.415 |
| Peddocks | MigRate−1e−3; MigTime−1e−3 | 261092.252 | 415074.775 |
| Peddocks | MigRate−1e−4; MigTime−1e−0 | 7998.310 | 5129.392 |
| Peddocks | MigRate−1e−4; MigTime−1e−1 | 38127.645 | 37175.244 |
| Peddocks | MigRate−1e−4; MigTime−1e−2 | 36668.791 | 44216.707 |
| Peddocks | MigRate−1e−4; MigTime−1e−3 | 20731.241 | 87879.816 |
| Peddocks | MigRate−1e−5; MigTime−1e−0 | 7649.399 | 4681.135 |
| Peddocks | MigRate−1e−5; MigTime−1e−1 | 5427.468 | 3352.474 |
| Peddocks | MigRate−1e−5; MigTime−1e−2 | 13939.171 | 34187.621 |
| Peddocks | MigRate−1e−5; MigTime−1e−3 | 3125.211 | 1511.769 |

**Figure S13: Impact of simulated migration on the uniformity of individual singleton contributions.** Chi-squared test statistics measuring the deviation of observed individual singleton counts in the Bumpkin Island (A) and Peddocks Island (B) samples from the null hypothesis that each mouse contributes the same number of singletons. (Left) Magnitude of the chi-squared test statistic computed for each simulated (gray and multicolored) and empirical (black) chromosome. (Right) Corresponding summary of the mean and standard deviation of the chi-squared test statistics compute across chromosomes derived from the empirical data (black), simulated data without migration (gray), and simulated data with migration (multicolored). "Params" column denotes the parameter values used in the migration simulations, where "MigRate" denotes the rate of migration between the focal population and unsampled "ghost" population and "MigTime" denotes the proportion of the split time over which continuous, symmetric migration occurs into the present. For clarity, we only show the simulation results derived from the Bumpkin-World's End demographic model (for A) and the Peddocks-World's End demographic model (for B), though we note that patterns are similar regardless of which specific demographic model is used for a given population.

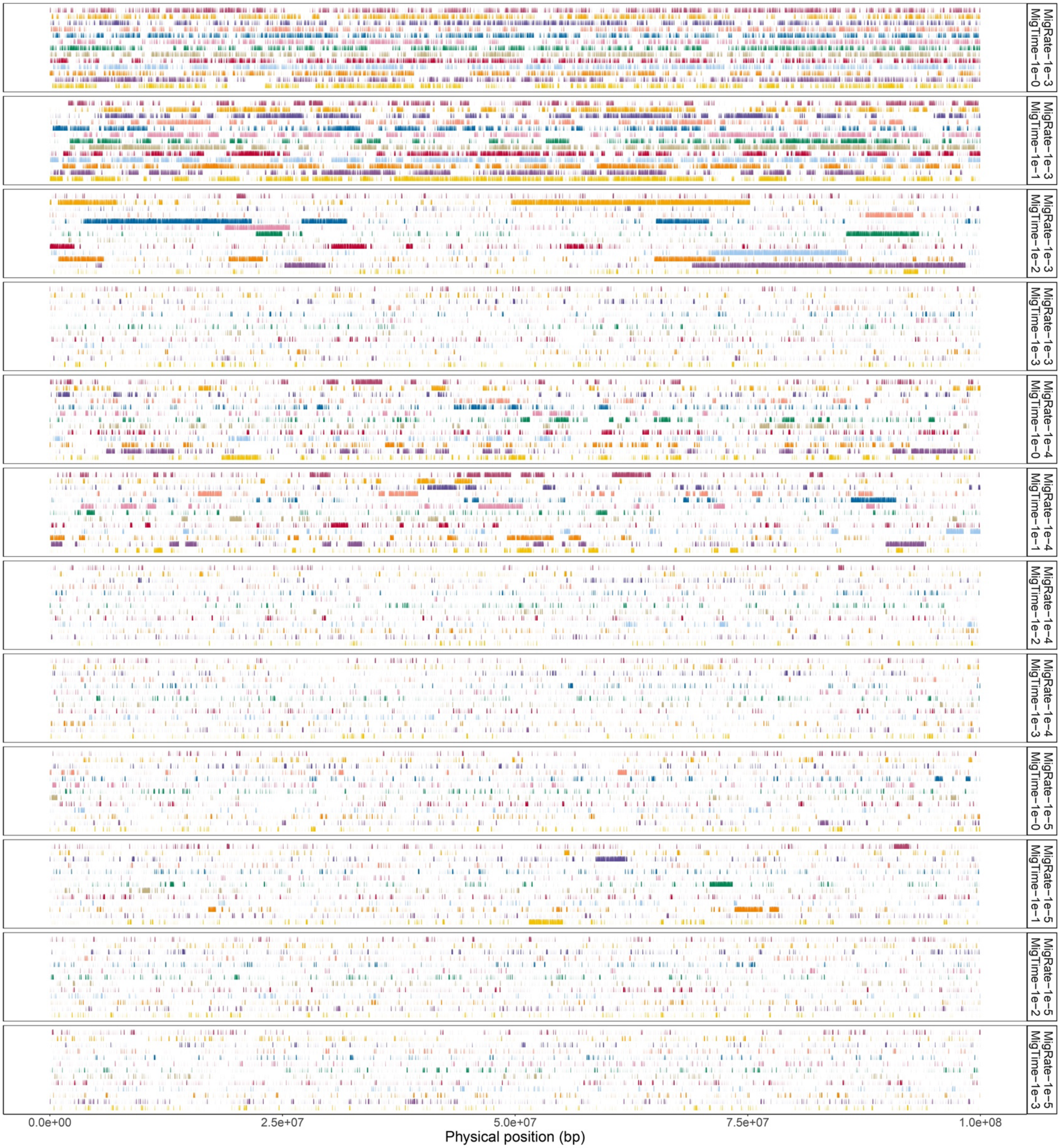


**Figure S14: Genomic distribution of singleton variants in the Bumpkin Island sample under simulated migration.** X-axes give the physical position along a representative, simulated 100

Mbp chromosome. Vertical, colored tick marks denote observed singleton variants. Each row reflects the singleton variants of a distinct simulated individual. Plots are faceted according to simulated migration regime where “MigRate” denotes the rate of migration between Bumpkin Island and the unsampled “ghost” population and “MigTime” denotes the proportion of the split time over which continuous, symmetric migration occurs into the present. For clarity, we only show the simulated singleton maps derived from the Bumpkin-World’s End demographic model, though we note that patterns are similar regardless of which specific demographic model is used for the Bumpkin Island sample.

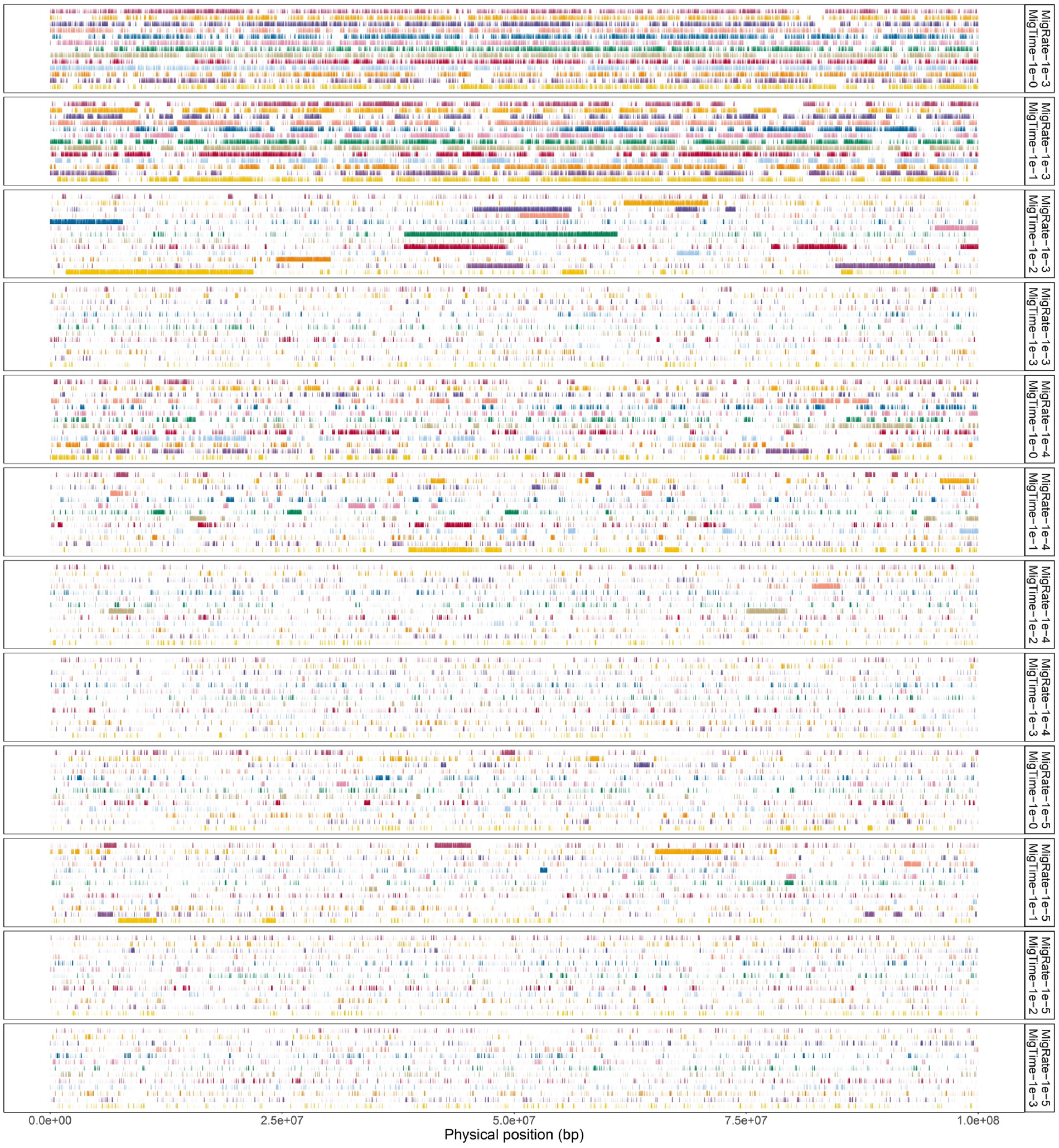


**Figure S15: Genomic distribution of singleton variants in the Peddocks Island sample under simulated migration.** X-axes give the physical position along a representative, simulated 100

Mbp chromosome. Vertical, colored tick marks denote observed singleton variants. Each row reflects the singleton variants of a distinct simulated individual. Plots are faceted according to simulated migration regime where “MigRate” denotes the rate of migration between Peddocks Island and the unsampled “ghost” population and “MigTime” denotes the proportion of the split time over which continuous, symmetric migration occurs into the present. For clarity, we only show the simulated singleton maps derived from the Peddocks-World’s End demographic model, though we note that patterns are similar regardless of which specific demographic model is used for the Peddocks Island sample.

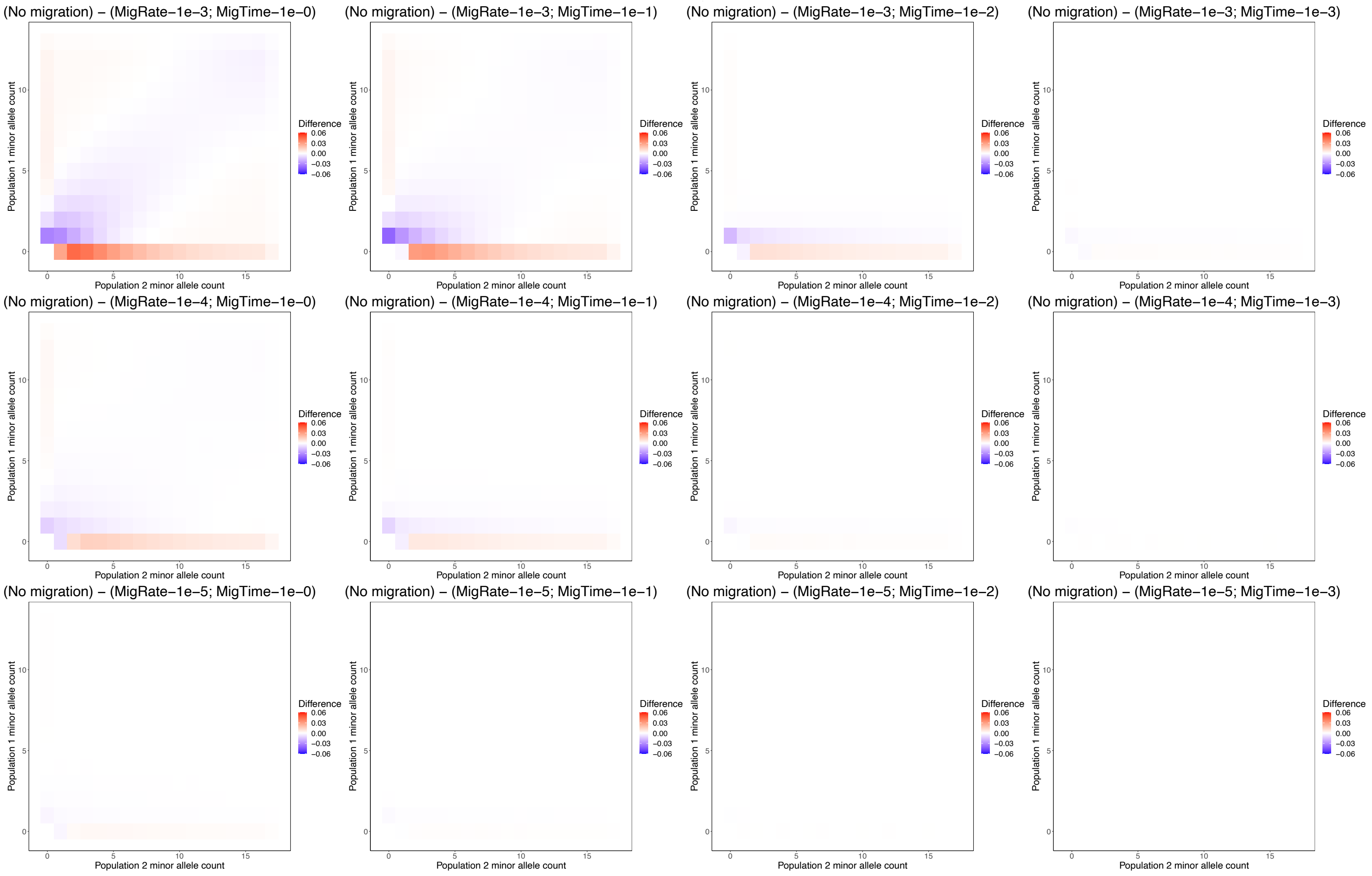


**Figure S16. Impact of ghost migration on the jSFS between Bumpkin Island and mainland World's End.** Each heatmap measures the impact of a distinct simulated migration regime on the proportion of SNPs in each bin of the jSFS between Bumpkin Island and mainland World's End. Within a heatmap, each cell represents the difference in the proportion of SNPs between simulations with and without migration (calculated as $x_{ij}^{no\ migration} - x_{ij}^{migration}$, where $x_{ij}$ represents the proportion of total SNPs that have a minor allele count of $i$ in mainland World's End (x-axis) and $j$ in Bumpkin Island (y-axis)). Warmer colors indicate that migration decreases the proportion of SNPs in a given jSFS bin, while cooler colors indicate that migration increases the proportion of SNPs. Plot titles denote the parameter values used in the migration simulations, where "MigRate" represents the rate of migration between the focal population and unsampled ghost population and "MigTime" represents the proportion of the split time over which continuous, symmetric migration occurs into the present. Plots are arranged such that the highest intensity migration regime is depicted in the top left, while the lowest intensity regime is depicted in the bottom right. Across plots, the duration of migration ("MigTime" parameter) decreases from left to right and the rate of migration ("MigRate" parameter) decreases from top to bottom.

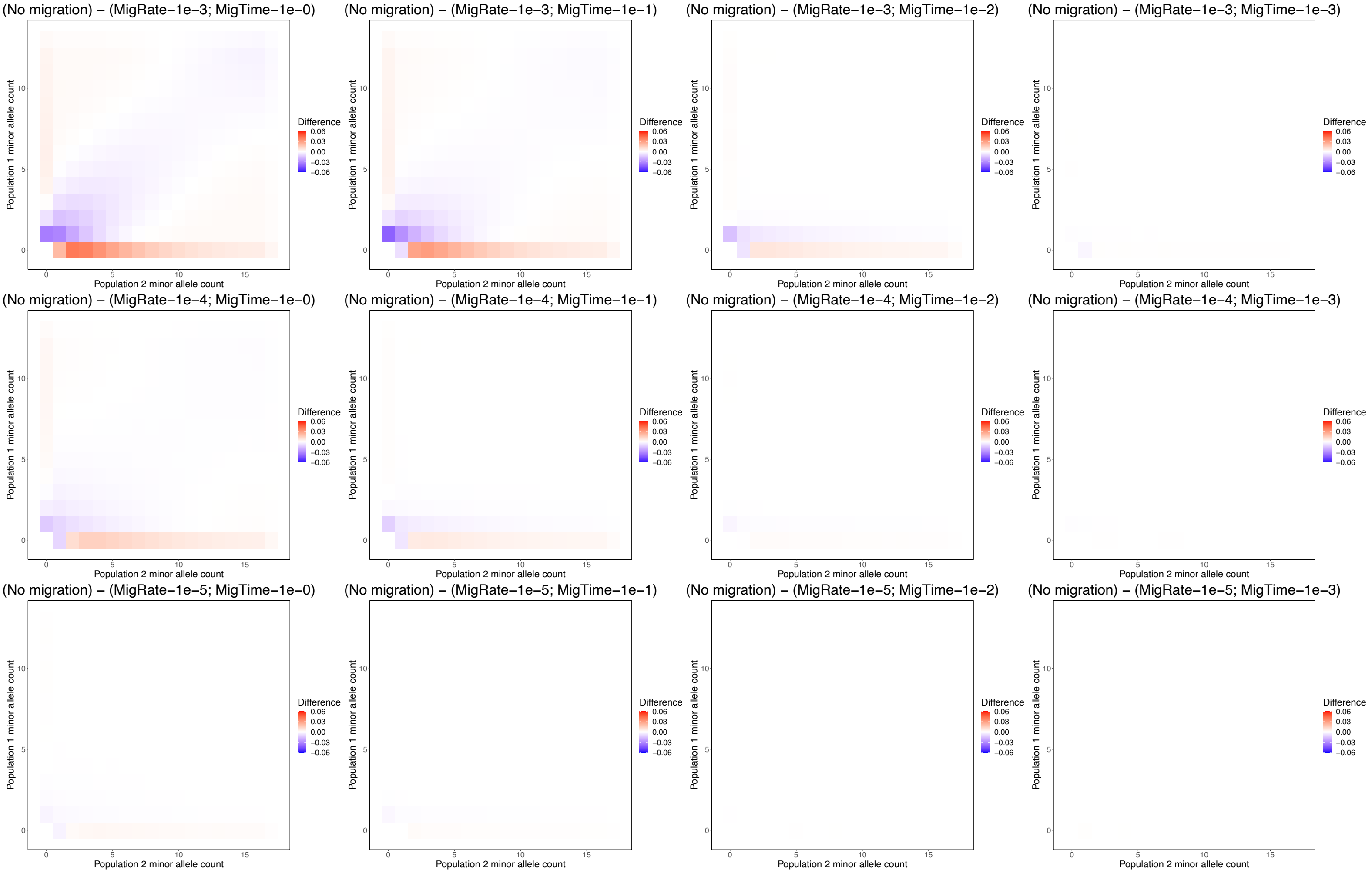


**Figure S17. Impact of ghost migration on the jSFS between Peddocks Island and mainland World's End.** Each heatmap measures the impact of a distinct simulated migration regime on the proportion of SNPs in each bin of the jSFS between Peddocks Island and mainland World's End. Within a heatmap, each cell represents the difference in the proportion of SNPs between simulations with and without migration (calculated as $x_{ij}^{no\ migration} - x_{ij}^{migration}$, where $x_{ij}$ represents the proportion of total SNPs that have a minor allele count of $i$ in mainland World's End (x-axis) and $j$ in Peddocks Island (y-axis)). Warmer colors indicate that migration decreases the proportion of SNPs in a given jSFS bin, while cooler colors indicate that migration increases the proportion of SNPs. Plot titles denote the parameter values used in the migration simulations, where "MigRate" represents the rate of migration between the focal population and unsampled ghost population and "MigTime" represents the proportion of the split time over which continuous, symmetric migration occurs into the present. Plots are arranged such that the highest intensity migration regime is depicted in the top left, while the lowest intensity regime is depicted in the bottom right. Across plots, the duration of migration ("MigTime" parameter) decreases from left to right and the rate of migration ("MigRate" parameter) decreases from top to bottom.
